# A helminth mimic of TGF-β, TGM, enhances regenerative cutaneous wound healing and modulates immune cell recruitment and activation

**DOI:** 10.1101/2022.09.24.509317

**Authors:** Katherine E. Lothstein, Fei Chen, Pankaj Mishra, Danielle J. Smyth, Wenhui Wu, Alexander Lemenze, Yosuke Kumamoto, Rick M. Maizels, William C. Gause

**Affiliations:** Center for Immunity and Inflammation, Department of Medicine, New Jersey Medical School, Rutgers, The State University of New Jersey; Newark, NJ 07103, USA; Center for Immunity and Inflammation, Department of Pathology, Immunology, and Laboratory Medicine, New Jersey Medical School, Rutgers, The State University of New Jersey; Newark, NJ 07103, USA; Wellcome Centre for Integrative Parasitology, School of Infection and Immunity, University of Glasgow; Glasgow, UK

## Abstract

Intestinal helminth parasites express excretory/secretory (ES) molecules, which modulate the type-2 immune response including anti-inflammatory and tissue repair pathways. TGF-β mimic (TGM), an ES molecule secreted by *Heligmosomoides polygyrus* (Hp), binds TGF-β receptors yet lacks structural homology to TGF-β and exhibits distinct receptor interactions. We demonstrate TGM treatment enhanced wound healing and tissue regeneration in an *in vivo* wound biopsy model. TGM, in a 1.5% carboxymethylcellulose solution, was topically administered beneath a Tegaderm layer. Through histological analysis, increased restoration of normal tissue structure in the wound beds of TGM-treated mice was observed during mid- to late-stage wound healing. These observations included accelerated re-epithelialization and hair follicle regeneration, without increased scarring. Flow cytometric and gene expression analysis showed differential expansion of myeloid populations at different stages of wound healing. This included enhanced early accumulation and persistence of macrophages in TGM-treated wounds during the initial inflammatory phase. Additionally, the percentage of alternatively activated (M2) macrophages expressing CD206 was reduced with TGM treatment during early and mid-stage wound healing. scRNAseq analysis of TGM-treated wounds indicate upregulation of multiple wound healing-associated genes without expression of CD206 within macrophage subsets. Experiments with truncated TGM constructs revealed that the TGFβ-R binding domain was essential in enhancing the wound healing response. In summary, TGM can accelerate skin wound healing and pro-restorative maturation through its interaction with the TGF-β receptor and stimulate the recruitment and reprogramming of specific macrophage subsets. This study indicates a role for TGM as a potential novel therapeutic option for enhanced wound healing.

**One-Sentence Summary:** A helminth-derived protein leads to rapid wound closure, skin regeneration, and reprogramming of macrophage activation through TGF-βR binding.

## INTRODUCTION

Skin is an important barrier and defense mechanism in mammalian hosts and after an injury, the host must quickly respond with the activation of the wound healing response (*1-3*). Wound repair is a highly regulated system that clears the injured area of external pathogens and quickly covers the exposed area to prevent further damage and reduce infection (*1, 4*). This tissue repair process involves three main overlapping stages for tissue resolution: inflammation, proliferation, and maturation. The inflammatory stage involves the rapid recruitment of multiple cell types to the area of insult immediately after damage. The proliferation phase involves the activation of endothelial cells, macrophages, and fibroblasts that cover the wound. Finally, the maturation phase involves the differentiation and deposition of extracellular matrix (*1-5*).

The tissue repair response must be tightly regulated since delays or impairment of the process can also result in chronic injury or excessive fibrosis, which can limit subsequent function of the damaged tissue (*6-9*). Fibrosis is the prevailing consequence of wound repair that results in scarring in place of regeneration (*2*) and is considered a significant public health burden, resulting in billions of dollars spent each year (*6, 8, 10*). Therefore, the development of a treatment that both rapidly suppresses harmful inflammatory responses and induces the expression of tissue repair factors that favor tissue regeneration over scarring could promote more effective wound healing and lead to better clinical outcomes (*2*). The type 2 immune response includes both pro- and anti-inflammatory functions and is beneficial for resolving cutaneous wounds, as many type 2 molecules and signaling pathways also appear to have a role in the activation and upregulation of tissue repair factors (*2, 4, 11, 12*). However, excessive or uncontrolled inflammation is detrimental (*7, 8, 13*). Yet, studies have found that the type 2 immune response initiated through helminth infections has evolved specific mechanisms which prevent this dysregulated immune response (*12, 14, 15*). This is due to the existence of helminth-associated mechanisms that activate a more regulated type 2 immune environment to control the infection and limit the damage to the host (*16, 17*).

In previous studies, we demonstrated that during a mouse helminth infection, lung inflammation and acute lung injury were mitigated through IL-4R-dependent mechanisms that involved both control of harmful inflammation and activation of factors that directly enhanced tissue repair (*12*). Indeed, many of the same cells involved with the helminth-modified type 2 immune response play a role in initiating or augmenting the tissue repair response, including Th2 cells, eosinophils, ILC2s, and most importantly, alternatively activated (M2) macrophages (*7, 11, 18*). Yet, helminths themselves produce comorbidities, raising concerns regarding their use as well as practicality for therapeutics. Recent studies, however, have shown that helminths produce specific factors that can modulate the host type 2 immune response (*19, 20*). These excretory/secretory (ES) molecules comprise a number of distinct proteins, carbohydrates, and lipids (*21, 22*). One, in particular, is referred at TGF-β mimic (TGM), as it can bind the TGF-β receptor, and was originally isolated from an ES supernatant obtained from cultures with the intestinal nematode parasite, *Heligmosomoides polygyrus*. TGM, while sharing no sequence similarity to the TGF-β family, can bind to host TGF-β receptors in both humans and mice (23), ligating the two receptor subunits through the independent N-terminal 3 domains of the protein (24). TGM can induce Foxp3+ regulatory T cells *in vitro* to an equal or greater extent than TGF-β (*25, 26*) and exerts anti-inflammatory effects *in vivo* in models of colitis (*27*) and airway allergy (*28*). In this work, we investigated whether TGM administration can modulate the progression of wound healing in a previously described cutaneous wound biopsy model. Our studies showed that when administered under a Tegaderm bandage, TGM markedly enhances wound closure, re-epithelialization, collagen crosslinking, and stimulates hair follicle formation, thereby promoting tissue regeneration over tissue scarring.

## RESULTS

### Topical TGM treatment accelerated wound closure

Previous studies have suggested helminth infections can trigger immunomodulatory and wound healing effects on surrounding tissues and that associated helminth excretory/secretory (ES) products may contribute to this response (*12, 17*). To examine specifically whether ES products might promote tissue repair, the effects of ES products from *Heligmosomoides polygyrus* (HES) were analyzed using an *in vitro* scratch test. In this standard test, HES was added to a culture including a 50:50 mix of an immortalized human keratinocyte cell line (HaCaT) and a mouse fibroblast cell line (L929), as previously described (*29, 30*). This test demonstrated an enhanced cell migration by HES compared to media alone (Fig. 1a, b). The recent identification of a purified recombinant HES product, TGM, that has an apparent immunomodulatory role (*15, 23*) and the ability of TGM to bind the TGF-βR (receptor), which has previously been shown to be important in wound healing (*31, 32*), raised the possibility that TGM may contribute to the effects of HES. TGM was thus employed in the scratch test used to assess HES activities. As shown in Fig. S1a, administration of TGM alone showed enhanced wound closure over a 24-hour time interval.

**Fig 1.**
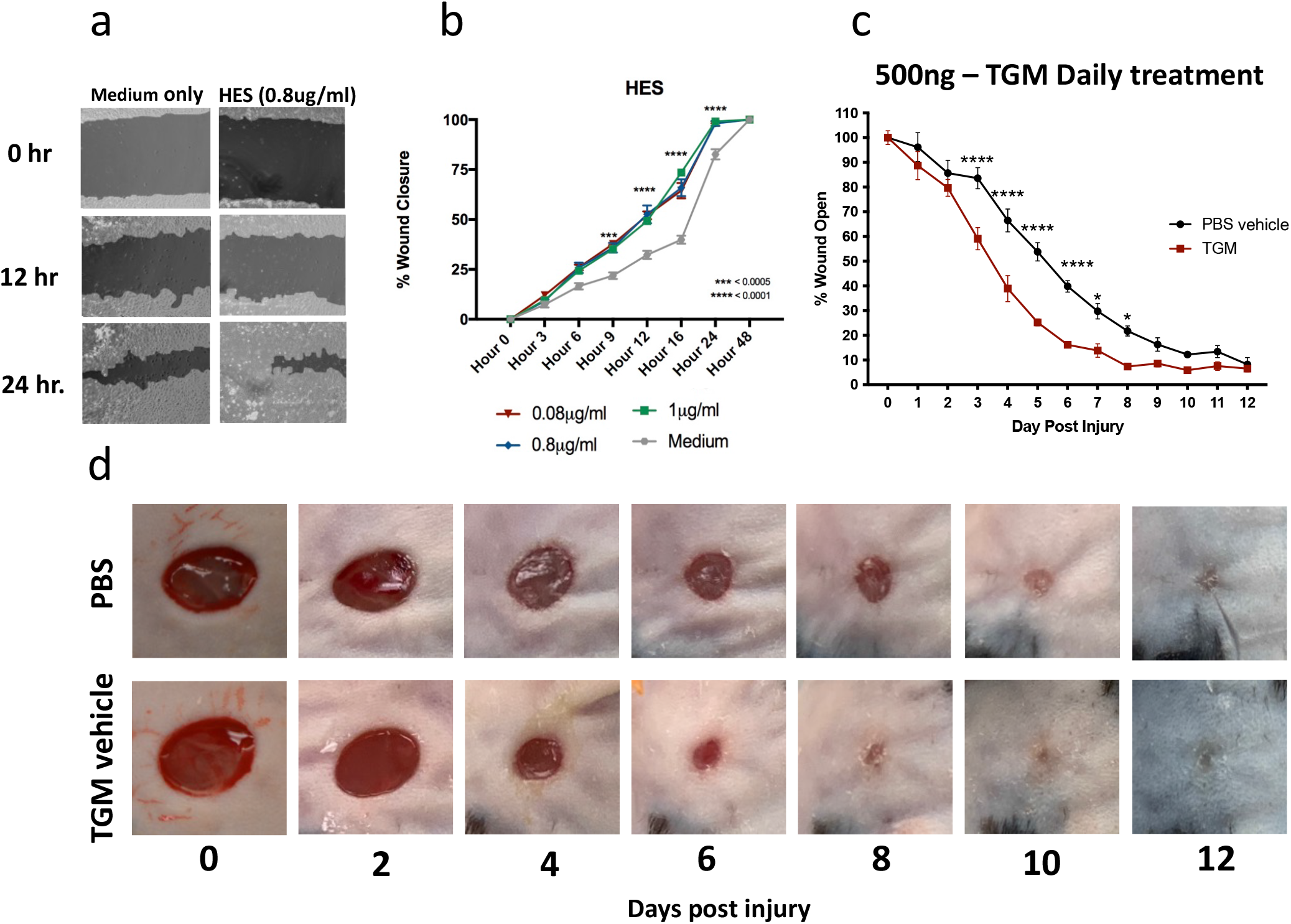
HES and the purified TGF-β mimic, TGM, accelerates wound closure in both a 2-D scratch test and murine dorsal wounds. (**A, B**) A 2-D *in vitro* scratch test wound model was generated with a 50:50 co-culture of L929 fibroblast and HaCaT keratinocytes to examine the wound closure and migration with the application of HES. (A) Representative images, produced from the analysis through the online program, Tscratch, identify the area of open wounds for quantitation (filled-in areas). (B) The area of wound remaining open, calculated by Tscratch, was used to quantify the rate of wound closure for different concentrations of HES (0.08 – 1ug/ml) compared to media alone from 0 to 24 hours. The percentage of wound closure at each time point is compared to the percentage open at hour 0 (** p <0.01, *** p < 0.001, **** p < 0.0001; n = 5 independent wells per condition). (**C**) 5mm full-thickness excisional wounds were generated on the dorsal skin of C57BL/6 mice. Wounds were treated with PBS vehicle control or TGM (500ng) and covered with Tegaderm^©^ for the duration of the study. Wound size analysis (ImageJ) was performed on the gross images obtained at each time point during the course of treatment. Treatments were given daily, while the dressing was changed every other day. Wound closure rates with either a daily dose of topical 500ng TGM or PBS vehicle control over 12 days were quantified as the percentage of wound closure at each time point compared to the percentage open at Day 0. Results from two or more independent determinations demonstrated similar results (** p <0.01, *** p < 0.001, **** p < 0.0001; 5 independent wounds from each treatment were measured through blinded analysis on ImageJ). (**D**) Representative wound images of mice treated topically with PBS vehicle control or TGM (500ng) on days 0, 2, 4, 6, 8, 10, and 12 demonstrate the rate of wound closure over the course of treatment. Statistical analysis was performed using a two–way analysis of variance (ANOVA) test (B, C) with Tukey’s multiple comparisons for comparison between all treatment groups at each timepoint. Error bars represent mean+/- SEM.

Given the enhanced closure in the *in vitro* wound healing model, we next examined the effects of TGM in a standard murine full-thickness skin wound biopsy model, as previously described (*33-35*). The presence in mice of an extensive subcutaneous striated muscle layer, the panniculus carnosus, reduces tension in wounds and activates wound contraction at a much faster rate than in humans (*36-38*). Several models have been developed that are more comparable to human skin, where re-epithelialization plays a more dominant role, that include various methods delaying wound closure including skin fixtures and Tegaderm (3M, St. Paul, MN) (*33-35*). Studies using Tegaderm alone have demonstrated similarly reduced rates of contraction as observed in sutured fixation methods (*34*), making them a good model for human skin wound healing and associated tissue remodeling. Consistent with previous studies (*33-35*), we found that our new protocol employing Tegaderm overlaying the wound provides both protection over the exposed wound and supplies enough tension around the wound to reduce excessive contraction and allow observation of re-epithelialization (Fig. S1b, c).

To further assess the therapeutic potential in this model, the TGM molecule was applied as a topical reagent, a common method of drug application for local wound treatment. A vehicle with 1.5% carboxymethylcellulose in PBS was used to generate a more viscous solution that would remain on the wound surface (*39*). TGM was mixed with the vehicle for application and was injected underneath the Tegaderm and on the surface of the wound after a 5mm biopsy punch was administered to the dorsal skin of the mice (n=5) (Fig. S1b, c). We initially tested various concentrations of TGM (1ng – 500ng/wound) as well as different treatment regimens ranging from one dose during wounding to one dose given every other day (Fig. S2a). After performing multiple dose-dependent studies at different concentrations and dosing schedules, we identified a daily dose of 500ng of TGM, applied to each wound, as the concentration that provides the most significant, reproducible results (Fig. 1c; Fig. S2b). As shown in Fig. 1 c and d, daily topical application of 500ng to a 5mm biopsy wound covered with Tegaderm resulted in significant improvement as early as day 3 with TGM leading to wound closure of 40% compared to day 0. In contrast, the PBS vehicle control group only obtained a 15% closure by that same time. The significant effect of daily TGM was maintained between days 4 and 10 with a sustained 30-40% difference in wound closure between the two groups at each time point. (Fig. 1c, d). By day 10, both treatment groups had closed to a degree in which differences could no longer visually be observed.

Concurrent with the enhanced wound closure, there was a noted increased wound exudate accumulating under the Tegaderm of the TGM-treated wounds, particularly at days 4 and 5 (Fig. S3a). This exudate was identified as serous discharge which is light-yellow in nature, less viscous than fluid normally associated with infection, and generally considered beneficial for wound healing (*40*). LC-MS/MS protein analysis of this liquid demonstrated a significant increase in haptoglobin and thrombospondin-4 in the TGM-treated wounds (Fig. S3b, c), which are known to influence macrophage differentiation and epithelial cell migration, respectively (*41-44*). Collectively these studies showed that TGM significantly enhances the rate of wound closure with increases in serous volume and concentrations of factors associated with augmented wound healing.

### TGM increased the granulation of tissue

The maturation phase of wound healing that occurs after wound closure includes the recruitment and activation of various immune cell populations that can influence whether the healing process favors a pro-fibrotic pathway associated with scarring or a more favorable pro-regenerative pathway (*45-47*). To examine whether TGM modulates later stages of wound healing resolution, histological analysis of the wound bed was performed after wound closure. Wound tissue granulation is a dermal tissue matrix that replaces the initial clot and fills the wound space with newly formed capillaries, epithelial cells, and infiltrating immune cells (*48, 49*). The wound bed tissue was analyzed for cellular infiltrate and extracellular matrix, indicators of tissue granulation. TGM-treated wounds showed accelerated granulation formation in comparison to PBS-vehicle control treated wounds. Increased granulation tissue was present as early as days 4 and 5 (Fig. 2a, b) and was associated with significantly greater wound bed thickness of the tissue by day 5 compared to the control (Fig. 2c). This enhanced thickness suggests that TGM is amplifying the formation and increasing the strength of the developing tissue as it heals and matures (*48, 49*).

**Figure. 2.**
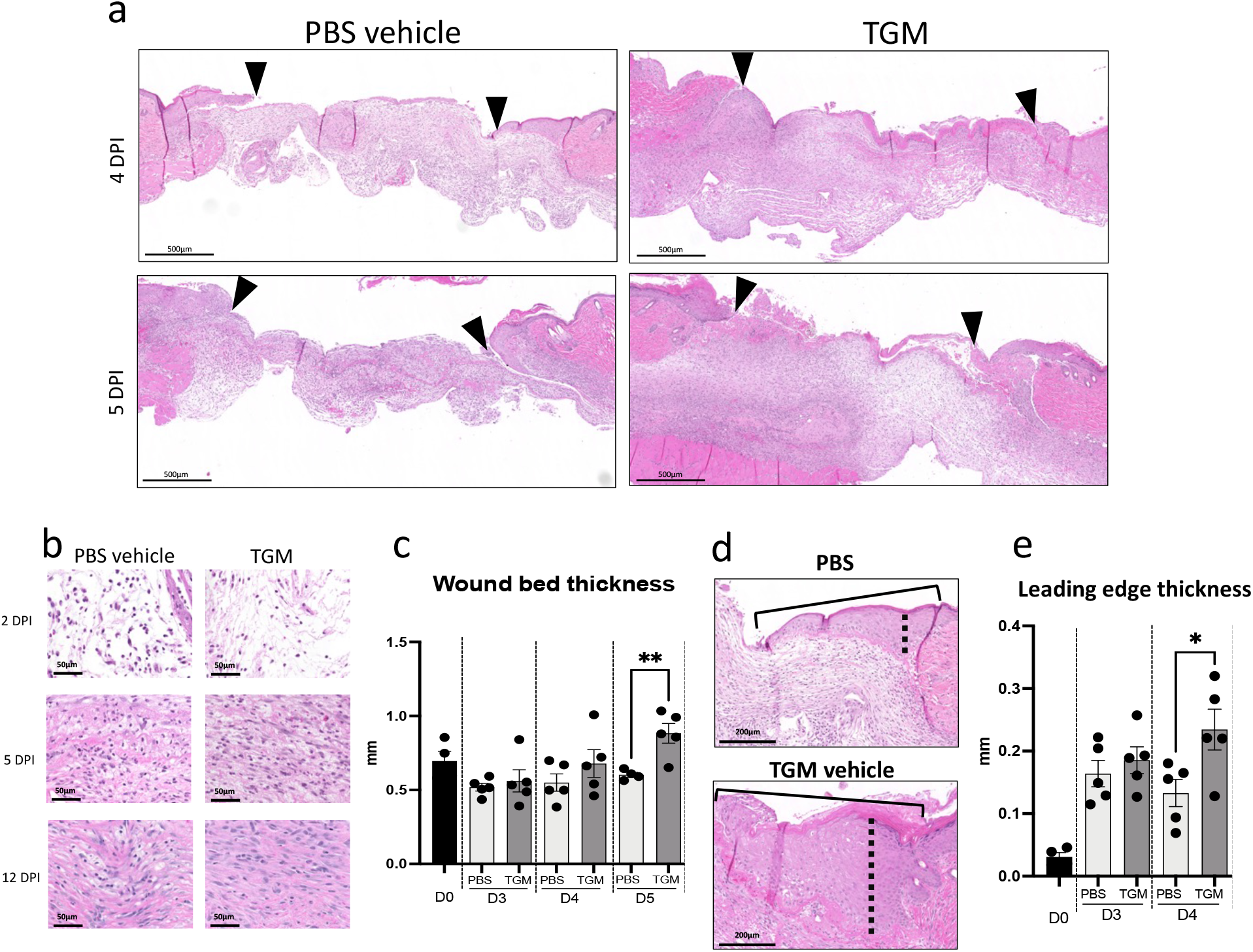
TGM enhances the granulation development and thickness of the epidermis/dermis within the newly formed wound. (**A**) Representative hematoxylin and eosin (H&E) stained images on days 4 and 5 post-injury. Eosin staining was used to identify granulation tissue formation within the wound beds (marked by arrows) of the TGM and PBS vehicle control wounds at each timepoint. (**B**) Representative H&E images at higher magnification (20x) were used to visually assess the cellular and extracellular matrix deposition within the wound beds of PBS vehicle control and TGM-treated wounds on days 2, 5, and 12 post-injury. (**C**) H&E stained wounds were used to measure the wound thickness of the treatment groups on days 3, 4, and 5 post-injury. Blinded analysis in ImageJ was used to measure the thickness of the granulation tissue within the wound bed (area between the black arrows in (A)). Multiple lengths obtained from one wound bed were averaged. Results from two or more independent determinations demonstrated similar results (** p <0.01; 5 biologically independent samples were used per treatment for the blinded analysis on ImageJ). (**D**) Representative H&E images of the wound edge, marked by the solid black line, were used to measure wound edge thickness. Blinded analysis in ImageJ was used to measure the thickness of the wound edge. The dotted black line indicates the length measured. (**E**) H&E stained images were used to measure the thickness of the leading edge within the wound bed. Measurements for PBS vehicle control and TGM were analyzed on days 0, 3, and 4. The thickest part of the wound edge over the granulation tissue was used for the analysis. Results of two or more independent determinations demonstrated similar results (**p <0.05; 5 biologically independent samples were used per treatment for the blinded analysis on ImageJ). Statistical analysis was performed using a t-test (C, E) to compare the two treatments at each time point. Error bars represent mean+/- SEM.

Concurrent with the granulation process, the wound edges thicken as keratinocytes are activated to proliferate and migrate over the newly formed dermis that develops from the granulation tissue. This process initiates re-epithelialization (*50-52*). With TGM treatment, there was a significant increase in wound leading-edge thickness by day 4, which correlates with the enhanced wound closure that was first observed at this early time point (Fig. 2d, e). However, while increased granulation and wound closure after TGM administration demonstrate a faster rate of tissue reorganization, these observations do not indicate whether the tissue is following a fibrotic or regenerative developmental trajectory. Therefore, further analyses of collagen orientation, fibroblast differentiation, and hair follicle development were used to determine the wound healing pathway followed with TGM application.

### TGM increased dermal collagen, reduced fibrosis, and increased hair follicle formation

Fibrotic development can influence collagen orientation as scar tissue is generally associated with tight parallel collagen bundle alignment, which reduces the skin’s elasticity, limiting the tissue’s normal function (*1, 10, 53*). In contrast, skin tissue collagen typically exhibits a more cross-linked basket-weave orientation providing greater flexibility to the tissue (*54*). Image analysis of picrosirius red-stained tissue sections, obtained from formalin-fixed skin biopsy tissues, was performed using the MRI Fibrosis Tool plugin on ImageJ (*55*). The intensity of the stain was quantitated to assess the relative percentage of collagen development in the wound, as previously described (*56*). TGM treatment of skin biopsies accelerated collagen deposition by day 7 compared to treatment with PBS vehicle control (Fig. 3a, c). By day 7, collagen compromised approximately 50% of the new tissue composition in the TGM-treated wounds, but only 40% of the PBS vehicle control treated skin (Fig. 3a, c). However, by day 12, both treatment groups exhibited approximately 60% collagen composition (Fig. 3a, b, c). Yet, while collagen composition was similar between the two treatments, the collagen orientation was markedly different. On day 12, in the TGM-treated wounds, collagen exhibited the distinct basket-weave morphology characteristic of normal, unwounded skin (Fig. 3d; Fig. S4a) (*9, 54*). In contrast, PBS vehicle control treated wounds exhibited characteristic scar tissue morphology composed of parallel collagen bundles (Fig. 3d).

**Figure. 3.**
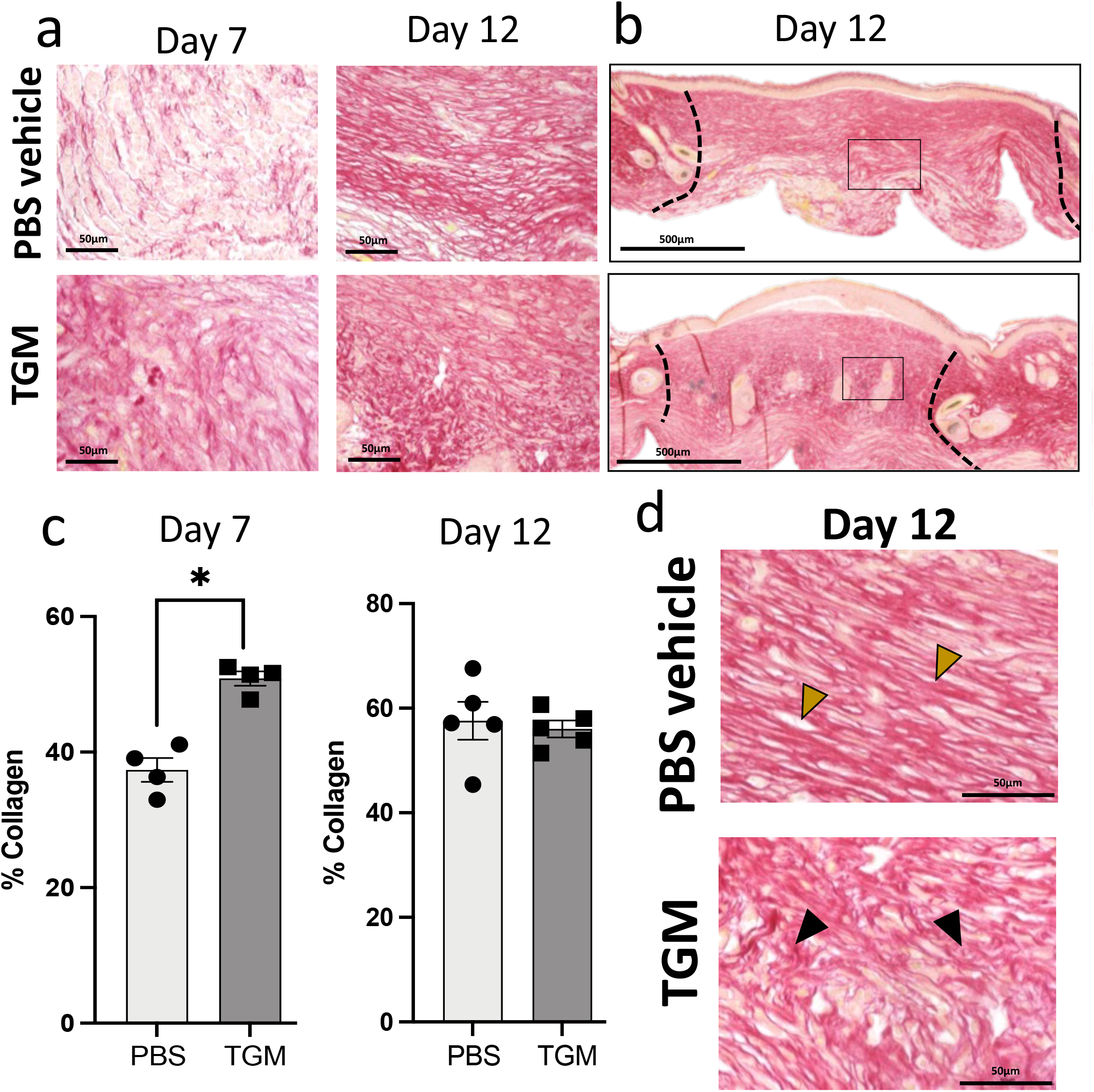
TGM-treated wounds are associated with pro-regenerative collagen deposition and orientation. (**A**) Picrosirius red stain was used to identify the orientation and quantify the percentage of collagen within the newly formed wound bed. Representative images of picrosirius red stain for collagen quantification (20x) identify the amount of collagen in the two treatments on days 7 and 12. (**B**) Two representative picrosirius red-stained images represent the whole wound bed of PBS vehicle control and TGM-treated wounds on day 12. The black box denotes the magnified region represented in (A). The black dotted lines indicate the wound edges on day 12. (**C**) The collagen area within the 20x picrosirius red-stained slides was quantified using the MRI Fibrosis Tool plugin on ImageJ. Multiple images were taken for each sample spanning the length of the wound bed and then averaged for that sample. Results from two or more independent determinations demonstrated similar results (** p <0.01; 5 biologically independent samples were used per treatment for the blinded analysis on ImageJ). Statistical analysis was performed using a t-test to compare the two treatments at each time point. Error bars represent mean+/- SEM. (**D**) Representative images of picrosirius red-stained slides on day 12 that were used to identify the collagen orientation in PBS vehicle control and TGM treated wounds. Yellow arrows highlight examples of collagen in a parallel orientation. Black arrows highlight examples of basket weave collagen deposition. Images represent similar images among the other 5 samples in each treatment.

A fibrotic skin tissue structure cannot support the neogenesis of hair follicles and hair growth. As a consequence, scarred skin typically lacks the presence of hair (*3, 46, 54*). Concurrent with the observation of enhanced basket weave morphology of collagen, we identified a significantly greater number of hair follicles at varying orientation and maturation levels in the newly formed dermis of TGM-treated wounds compared to the control wounds treated with the PBS vehicle control (Fig. 4 a, b). Epidermal extensions into the dermis, known as rete pegs, provide support and nutritional access to the multi-layers of the skin and are often used as a marker for tissue regeneration (*54, 57, 58*). Rete pegs were readily observed in TGM-treated but not PBS vehicle control treated wounds (Fig. 4b). At higher magnifications, the mature hair follicles were identified along with complete sebaceous glands and hair shafts, denoting normal skin development. (Fig. 4c). In contrast, in the PBS vehicle control treated wounds, the rete pegs were absent, and the dermal layer remained flat over the epidermis (Fig. 4d). Taken together, the increased development of hair follicles and rete pegs indicates enhanced skin tissue regeneration with reduced scarring following TGM treatment.

**Figure 4:**
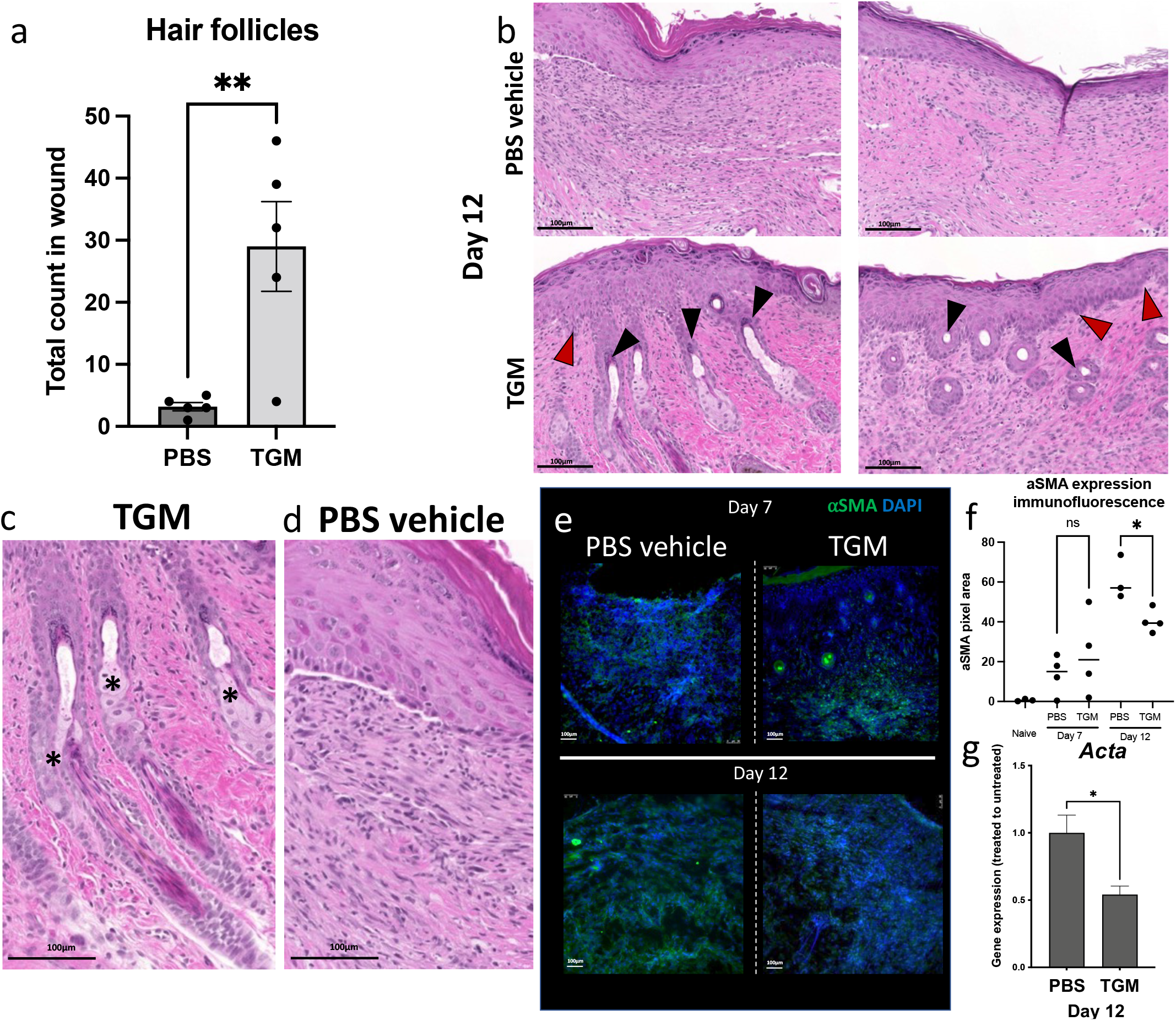
Enhanced hair follicle frequency coincides with regulated myofibroblast expression in TGM-treated wounds on day 12. (**A**) H&E stains were used to quantify the frequency of hair follicles within the wound beds on day 12. Hair follicles were included in the count if they were located within the wound bed and beneath the thickened epidermis which represented the area of the wound. (** p <0.01; 5 biologically independent samples were used per treatment for the blinded analysis on ImageJ). A t-test was performed to compare the two treatments at each time point. Error bars represent mean+/- SEM. Results from two or more independent determinations demonstrated similar results (**B**) Representative images of H&E stained slides from wounds on day 12 were used to identify skin maturation through the appearance of hair follicle formation and rete pegs within the wound bed. Black arrows = hair follicles, red arrows = rete pegs. (**C**) A representative magnified H&E image of a 12-day TGM treated wound shows the mature hair follicles, including sebaceous glands (*) within the wound bed compared to a representative image of a PBS vehicle control treated wound (**D**) that is lacking hair follicles and rete pegs (n = 5 for each condition and study). Images represent similar images among the other 5 samples in each treatment. (**E, F**) Alpha smooth muscle actin (αSMA) representing myofibroblast formation, was quantified in all images on days 7 and 12 through immunostained slides. (E) Representative immunofluorescent stained images of wound beds stained with αSMA on days 7 and 12; αSMA (green fluorescent signal), DAPI (blue fluorescent signal). (F) Immunostained slides with αSMA were quantified on days 7 and 12 by the fluorescent intensity measured as pixel area using ImageJ. (* p <0.05; 5 biologically independent samples were used per treatment for the blinded analysis on ImageJ). Results from two or more independent determinations demonstrated similar results (**G**) Gene expression analysis of *Acta2* expression was used to further analyze the αSMA expression within the wound bed on day 12 post-wound. (* p <0.05; 5 biologically independent samples were used per treatment for the blinded analysis on ImageJ). Results from two or more independent determinations demonstrated similar results. Statistical analysis was performed using a t-test (F, G) to compare the two treatments at each time point. Error bars represent mean+/- SEM.

α-Smooth muscle actin (αSMA) expression is another marker used to characterize wound regeneration as it represents myofibroblast activation and differentiation which is critical for wound healing (*54, 59, 60*). However, excessive myofibroblast activation, which can be quantified through αSMA expression, is indicative of scar tissue development (*54, 59, 60*). To assess αSMA expression, skin tissue samples were frozen in OCT on days 7 and 12 after skin biopsy and then stained with an αSMA fluorescent marker to analyze expression within the wound bed. ImageJ was used to analyze the fluorescent intensity of the immunofluorescent stained slides to quantify the αSMA expression. Through blinded analysis, we determined that αSMA expression increased after day 0 and was similar in both groups on day 7 (Fig. 4e, f; Fig. S4b). However, by day 12, elevations in αSMA were significantly less marked in the TGM-treated wounds compared to the PBS vehicle control (Fig. 4e, f). Further gene expression analysis by qPCR of *Acta* (αSMA) levels confirmed attenuated expression of myofibroblasts in TGM-treated wounds on day 12 (Fig. 4g) (*60*). Together, these data suggest suppression of myofibroblast activation and reduced scarring in wound beds after TGM treatment. Collectively, these histological observations indicate that daily TGM application, in addition to accelerating wound closure, enhances pro-regenerative relative to pro-fibrotic tissue wound healing.

### TGM enhances wound healing and tissue repair through TGF-β receptor binding activity

TGF-β plays a role in multiple mechanisms associated with wound healing, which are initiated through binding to the heterodimer of TGF-βRI and II (*1, 32, 61, 62*). Since TGM also binds to this receptor complex, we tested the possible role of wound healing modulation through the specific TGF-βR binding activity of TGM. TGM is composed of 5 domains, with domains 1-3 (TGM D1-D3) containing the TGF-βR binding activity domains and domains 4 and 5 (TGM D4-D5) likely involved with additional, yet currently unknown, binding activities (Fig. 5a) (*24, 63*). Wound biopsies were treated daily with 500 ng of truncated recombinant TGM constructs of TGM D1-D3, or TGM D4-D5, as well as TGM D1-D5 (full-length TGM), and PBS vehicle control for 7 days (5 mice/treatment group). The gross image area measurement, compared to day 0, at each time point, demonstrated that TGM D1-D5 (500ng, n=5), as seen in our earlier experiments, led to significant wound closure compared to the PBS vehicle control (n=5) on day 4 with 50% of the wound closed compared to the 30% closed in the control. TGM D1-D3 (500ng, n=5) showed almost identical wound closure activity compared to TGM D1-D5 (Fig. 5b, c). However, TGM D4-D5 (500ng, n=5) did not show enhanced wound healing relative to the PBS vehicle control treatment (Fig. 5b, c). Both TGM D1-D5 and TGM D1-D3 showed significant wound closure compared to TGM D4-D5 as early as days 3 and 4. These findings demonstrate that the wound healing activity of TGM is mediated specifically through TGF-βR binding.

**Figure. 5:**
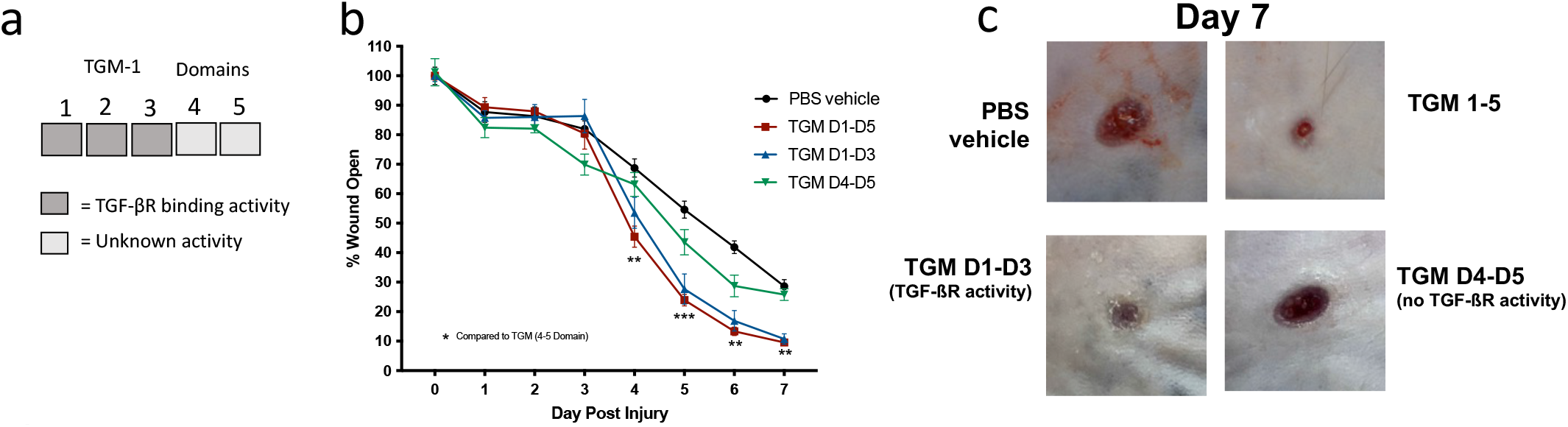
*in vivo* wound biopsy with truncated variants suggest TGM enhances wound healing through TGF-ßR activity. (**A**) The TGM molecule contains 5 domains. Domains 1 through 3 comprise the TGF-ßR domain activity. The activity of domains 4 and 5 is currently unknown. (**B**) 5mm full-thickness excisional wounds were generated on the dorsal skin of C57/Bl6 mice. Wounds were treated with PBS vehicle control or whole TGM (TGM D1-D5; 500ng) or TGM containing only domains 1 through 3 (TGM D1-D3; 500ng) or domains 4 and 5 (TGM D4-D5; 500ng) and covered with Tegaderm^©^ for the duration of the study. Wound size analysis was performed on the gross images obtained at each time point during the course of treatment. Treatments were given daily while the dressing was changed every other day. Wound closure rate with either a daily dose of topical TGM D1-D5, TGM D1-D3, TGM D4-D5, or PBS vehicle control over 7 days was quantified as the percentage of wound closure at each time point compared to the percentage open at Day 0. (** p <0.01, *** p < 0.001, **** p < 0.0001; Black stars represent the significance of TGM D1-D5 compared to TGM D4-D5; 5 independent wounds from each treatment were measured through blinded analysis on ImageJ). Statistical analysis was performed using a two–way ANOVA with Tukey’s multiple comparisons for comparison between all treatment groups at each timepoint. Error bars represent mean+/- SEM. (**C**) Representative wound images of mice treated topically with PBS vehicle control, TGM D1-D5, TGM D1-D3, or TGM D4-D5 (500ng) on day 7 provide a visual comparison of the area of wound remaining open between the 4 different groups. Results from two or more independent determinations demonstrate similar results

### TGM treatment increased macrophages but reduced the proportion of CD206+ alternatively activated (M2) macrophages

Different subsets of macrophages are involved in different stages and processes of wound healing (*2, 11*). To investigate the potential effects of TGM on macrophage activation, flow cytometric analyses were performed on tissues at various time points after *in vivo* wound biopsy treatments. Wounds were collected from mice and digested to obtain a single-cell suspension, as previously described (*50*). In the PBS vehicle treated control group, the macrophage population (CD11b+ F4/80+ CD64+) as a percentage of all CD45+ cells, was increased at 1 to 2 days post-injury and peaked between 5 to 7 days (Fig. 6a,c, shown in beige), similar to previously published results (*64*). This population comprises the majority of immune cells, with a marked higher frequency than dendritic cells and neutrophils (Fig. 6a). The macrophages still comprised a majority of the leukocytes throughout the study; however, after TGM treatment, macrophages peaked earlier at day 3, consistent with increased wound closure at early timepoints (Fig. 6b, c). From days 2 to 5, macrophages comprised a greater percentage of CD45+ cells in TGM-treated mice relative to the mice treated with the PBS vehicle control (Fig. 6b, c). By day 7, the TGM-treated macrophage population decreased while their counterparts in PBS vehicle control treated tissues increased significantly (Fig. 6c).

**Figure. 6:**
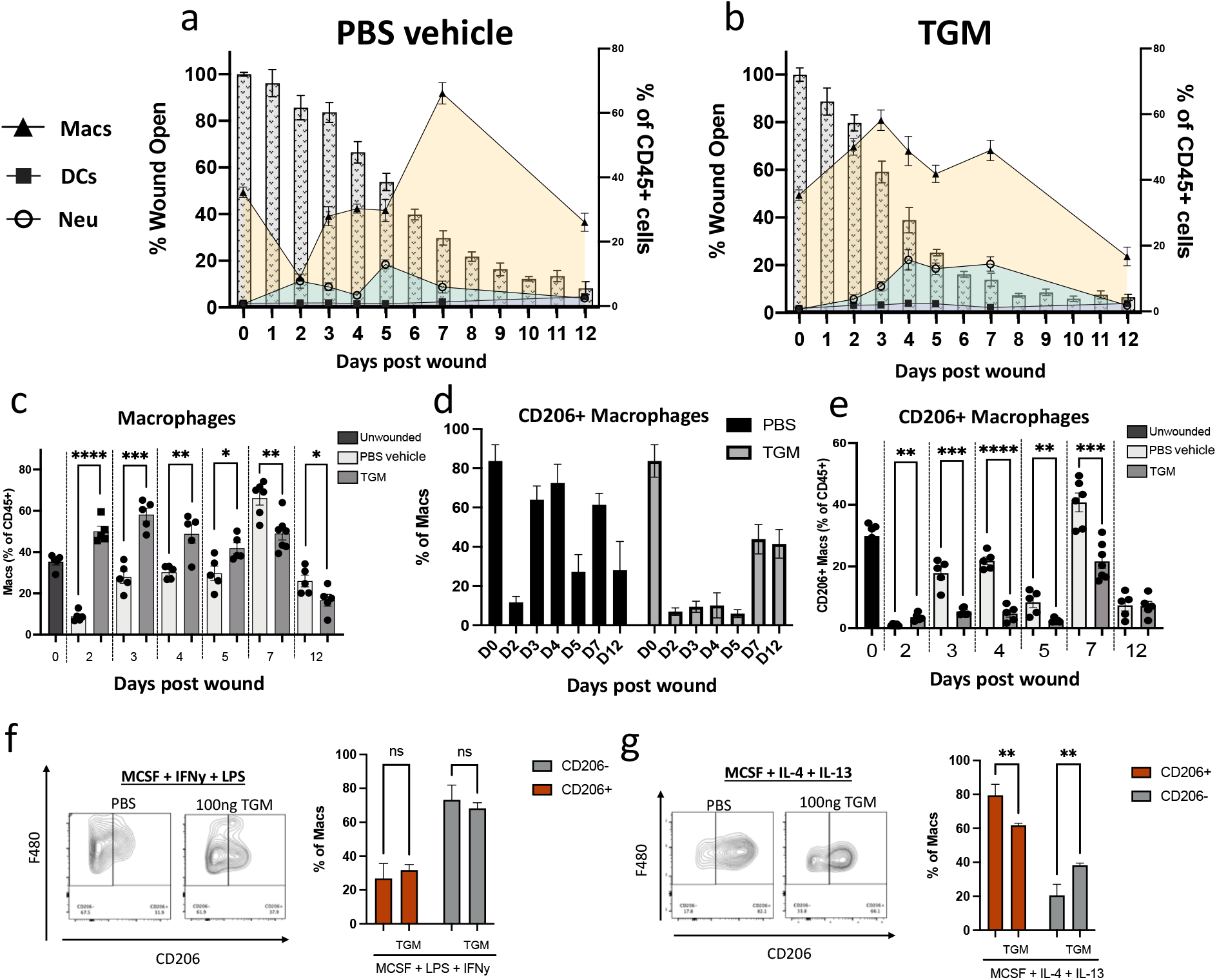
TGM treatment reprograms myeloid cell expansion and delays macrophage CD206 expression. (**A, B**) Flow cytometric analysis of the cell population within the wound beds from day 0 to day 12 for PBS vehicle control (A) and TGM (B) treated wounds. The analyzed cells include macrophages (CD11b+ F480+ CD64+), dendritic cells (MHCII+ CD11c+ CD64-) and neutrophils (CD11b+ Ly6G+ CD64-) as a percentage of the total CD45+ cells. The cell populations at each time point are overlayed on a bar graph representing the wound closure of each treatment group. Left Y-axis denotes the percentage of myeloid cells of total CD45+ cells represented in the line graph. Right Y access denotes the percentage of wound open as represented in the background bar graph. The X-axis represents the days post wounding. (**C**) Flow cytometric analysis of the frequency of macrophages (CD11b+ F480+ CD64+), as a percentage of all CD45+ cells, in PBS vehicle control and TGM treated wounds from day 0 to day 12. (* p <0.05, ** p <0.01, *** p < 0.001, **** p < 0.0001; 5 biologically independent samples were used per treatment at each time). Representative results from two or more independent experiments are shown. (**D**) Flow cytometric analysis for the frequency of the CD206+ macrophages as a percentage of total macrophages between PBS vehicle control and TGM treated wounds. Representative results from two or more independent experiments are shown. (**E**) Flow cytometric analysis of the frequency of CD206+ macrophages as a percentage of total CD45+ cells in PBS vehicle control and TGM treated wounds from day 0 to day 12. (* p <0.05, ** p <0.01, *** p < 0.001, **** p < 0.0001; 5 biologically independent samples were used per treatment at each time point). Representative results from two or more independent experiments are shown. (**F, G**) Flow cytometric analysis of the frequency of CD206+ macrophages gated on CD11b+ F4/80+. BMDMs were isolated and stimulated for 16 hours in a (F) classically activated (LPS/IFNy) macrophage-inducing environment or (G) alternatively activated (IL-4/IL-13) macrophage environment with or without TGM. Bar graphs represent the frequency of CD206+ or CD206-macrophages as a percentage of all macrophages (* p <0.05, ** p <0.01, *** p < 0.001, **** p < 0.0001; 3 wells were used per treatment). Statistical analysis of the CD206+ or CD206-populations was performed using a one–way ANOVA test with Tukey’s multiple comparisons for comparison between all treatment groups. Error bars represent mean+/- SEM.

During the wound healing transition from the proliferative phase to the maturation phase, alternatively activated (M2) macrophages, expressing CD206, expand and are thought to play a crucial role in the maturation stage of the wound healing process (*64-66*), typically located at the leading edge of the wound during the early phase where they interact and communicate with keratinocytes (*67*). In naïve skin, CD206+ macrophages are abundant, representing approximately 80% of all tissue-resident macrophages, as demonstrated in our unwounded samples (Fig. 6d) (*64*). To test the effect of TGM on the frequency of this specific macrophage subset, CD206+ macrophages were assessed on days 2, 4, and 6 after punch biopsy. PBS vehicle control treated mice showed a significant decrease in the percentage of CD206+ macrophages (CD11b+ F4/80+ CD64+) by day 2 that was largely restored by day 3 (Fig. 6d), consistent with previous studies in other skin wounding models (*64, 68*). However, this trend was not observed with TGM treatment. Similar to the PBS vehicle control group, on day 2, the CD206+ population decreased in the TGM treated group, but as wound healing progressed, increases in the CD206+ macrophages were delayed until day 7 (Fig. 6d). In both groups, as a percentage of total CD45+ cells, there was a peak accumulation of CD206+ macrophages on day 7, which was still significantly reduced in the TGM-treated wounds (Fig. 6e).

### TGM may induce a CD206-macrophage population expressing characteristic markers associated with wound healing

Our studies indicated marked changes in macrophage phenotypes due to TGM application after skin wounding. The wound healing response generates a type 2 immune pathway that activates CD206+ M2 macrophages in the presence of IL-4 and IL-13 (*69, 70*). The type 1 immune-associated response (induced by LPS and IFNγ) generates classically-activated macrophages that generally lack the CD206 marker (*69, 70*). To test whether TGM directly affects macrophage activation, bone marrow-derived monocytes were isolated from 8-week-old C57BL/6J mice (n=3) and treated with DMEM: F12 LCM conditioned lymphocyte growth media for 6 days, and the adhered macrophages were washed and stimulated, for 16 hours, with M-CSF and IFNγ/LPS or IL-4/IL-13 (Fig. 6f, g). As expected, only a small percentage (20%) of the cells treated with IFNγ/LPS expressed CD206, and the addition of TGM did not significantly affect the expression of this marker (Fig. 6f). In contrast, culture with IL-4 and IL-13 resulted in a marked increase in CD206+ macrophages (Fig. 6g). However, unexpectedly, the application of TGM significantly reduced the expression of CD206+ macrophages (Fig. 6g). These results indicate that TGM can directly influence macrophage activation in the context of the specific cytokine environment.

The reduced frequency of CD206+ macrophages concurrent with the enhanced wound healing in TGM-treated wounds appeared contradictory with previous studies suggesting a role for this macrophage subset in tissue repair (*64*). To further analyze potential heterogeneity in the activation of immune cells in the wound bed following TGM treatment, we performed single-cell RNA sequencing (scRNAseq). Cell suspensions were prepared on day 3 after wounding when marked changes were detected in myeloid cell populations using flow cytometric analysis following TGM treatment. scRNAseq using 10X genomics was performed on the CD45+ purified cells from three biological replicates treated with PBS vehicle control or TGM and labeled with 3 unique TotalSeq Hashtag Antibody markers (Biolegend, San Diego, CA). Using the unique hashtags, a single run for each treatment with all three biological replicates was performed. Hashtags were then used to identify samples from 3 individual mice within each treatment group.

Analysis of the six samples using ImmGen and standard cellular identification markers (*71-75*) revealed ten distinct clusters that were composed primarily of macrophage or monocyte populations, which typically comprise a majority of the immune cells at this early timepoint after cutaneous wounding (*64*), as confirmed by our results (Fig 6a). There were also several other significant populations including dendritic cells, natural killer cells, and neutrophils as defined by the top five expressed genes in each cluster in addition to identification with other gene profiles previously used to define each population (Fig. 7a, b; Fig. S5a, b) (*71*), which were also observed by FACS analysis (Fig. 6a). The proportions of the macrophage clusters were similar in each group while TGM was associated in an expansion of the clusters identified as neutrophils and Langerhans cells (Clusters 3 and 7) and diminution of dendritic cells (Clusters 4, 8 and 9) (Fig. 7c). Clusters 0 through 3 represented the most populated cell subsets in both treatment groups from the analysis. Clusters 0, 1, 2, and 5 resembled macrophages with their increased expression of *Adgre1* (F4/80) and *Itgam* (CD11b) (Fig. 7d) while Cluster 3 was identified as neutrophils with the expression of characteristic neutrophil markers (*Csf3r, Pglyrp1, IL-1β, Retnlγ*) (Fig. S6a) (*71*).

**Figure 7.**
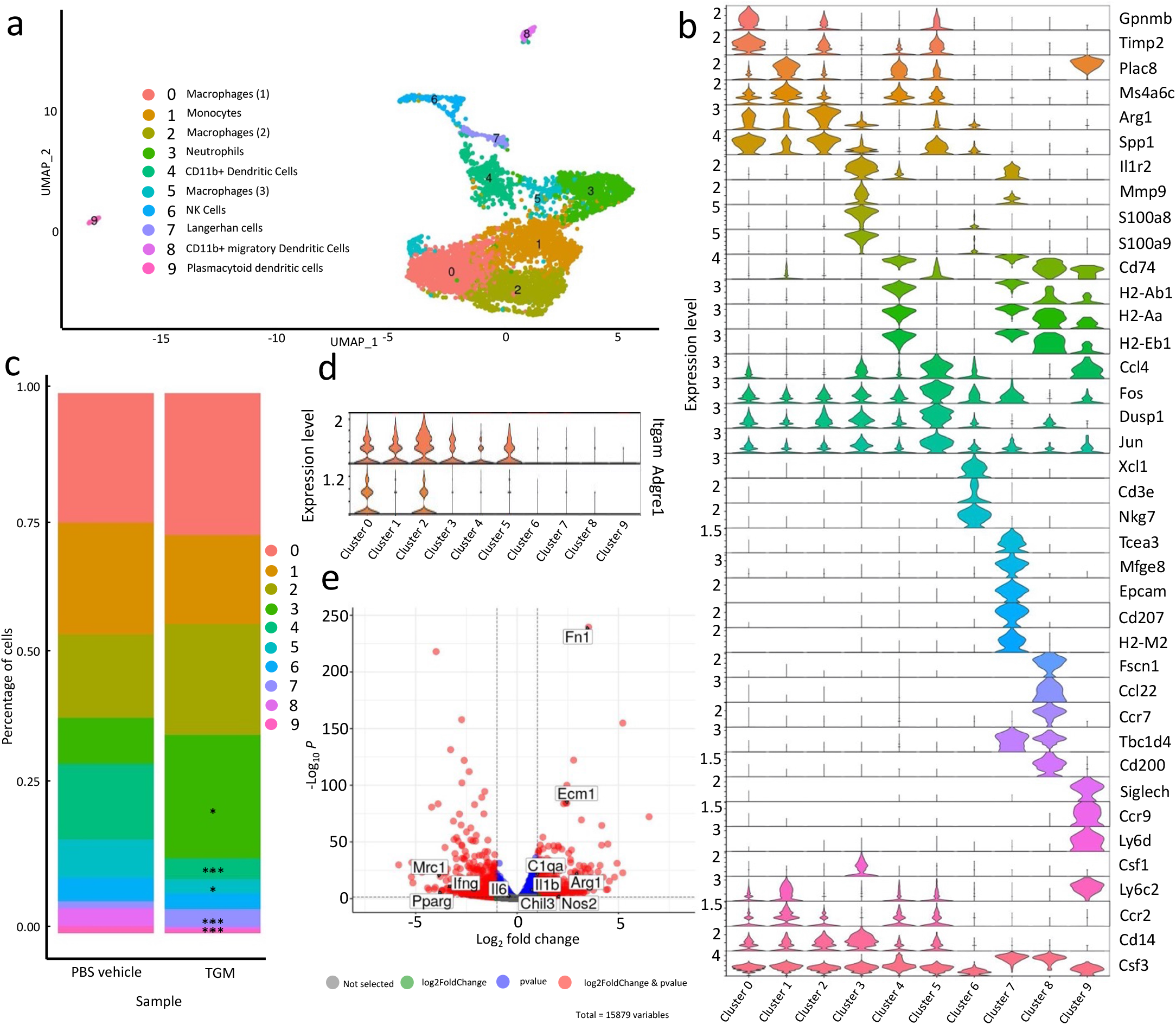
TGM treatment reprograms innate immune cell frequency and gene expression in cutaneous wound beds. scRNAseq analysis of CD45+ leukocytes obtained from wound bed at day 3 after skin injury. Skin wounding was performed as described in Fig. 1. (**A**) A uniform manifold approximation projection (UMAP) plot of the single cells obtained from CD45+ cells purified from PBS vehicle control and TGM treated wounds on day 3 after skin injury. 10X genomics scRNAseq using was performed on the CD45+ purified cells treated with PBS vehicle control or TGM (3 biologically independent samples were used per treatment labeled with 3 unique TotalSeq Hashtag Antibody markers). The analysis identified 9 distinct cell clusters. (**B**) Violin plots highlight the markers used to define the populations within the clusters. (**C**) The percentage of single-cell distribution among the clusters between the PBS vehicle control and TGM treated samples. Asterisks represent the significant difference in the cluster population size between the two treatment groups (* p <0.05, *** p < 0.001). (**D**) Violin plots represent the differential gene expression of macrophage markers (*Itgam* and *Adgre1*) within the clusters of TGM to PBS vehicle control combined. (**E**) Volcano plot representing the differential gene expression of the monocyte/macrophage populations between PBS vehicle control and TGM treated wounds (clusters 0, 1, 2, 5 combined). Horizontal dotted lines represent significant fold changes of expression for each treatment group. Vertical dotted lines represent a significant p value (<0.05). Red dots represent genes significantly upregulated within the two treatment groups.

To further dissect potentially more distinct macrophage subsets within these broadly defined clusters, we analyzed the differential gene expression of cells within the monocyte/macrophage clusters 0, 1, 2 and 5 between PBS vehicle control and TGM treated wounds. Using volcano plots, a considerable difference was detected in gene expression profiles between TGM- and PBS vehicle control treated cells within the defined macrophage clusters (Fig. 7e). Specifically, there was reduced expression of *Mrc1*, which encodes CD206 (*71*), in the TGM-treated compared to PBS vehicle control groups, while significant increases were observed in other M2 markers, including *Arg1* and *Fn1* (Fig. 7e, 8a) (*71, 76, 77*). Furthermore, another M2-associated marker, *Chil3*, while showing no difference in the combined macrophage populations between the two treatment groups (Fig. 7e), was increased in TGM-treated samples in clusters 0, 2, and 5 (Fig 8b, c). Notably, *Mrc1*/CD206 expression was abrogated in all macrophage clusters, with concomitant upregulation of other M2 products, suggesting a generalized loss of this marker rather than the outgrowth of a specialized CD206-negative subset. These findings suggest that wound healing-associated macrophages typically identified through *Arg1* and *Chil3* expression may not require CD206 expression to be identified as an M2 population.

**Figure 8.**
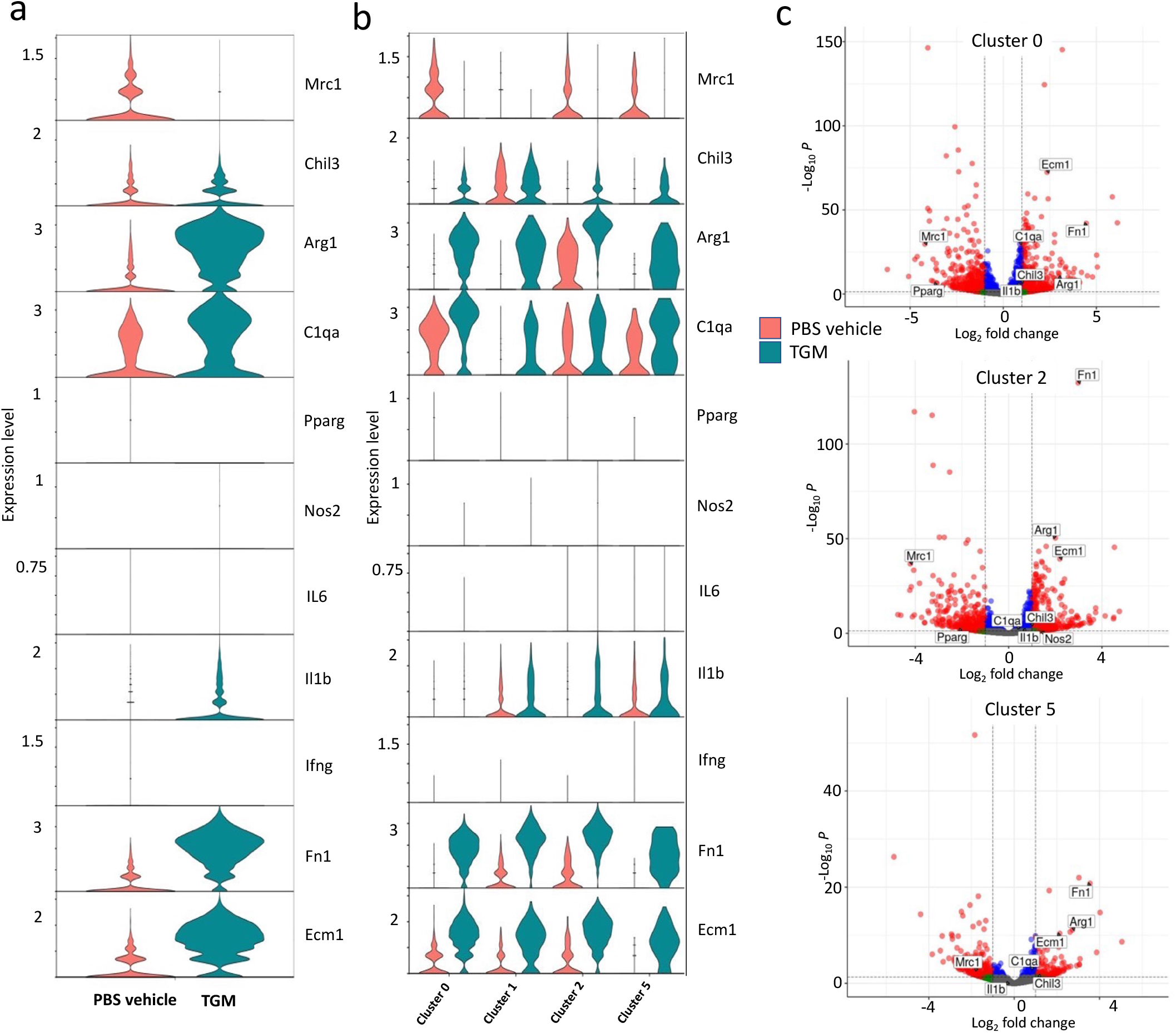
TGM-treated macrophages have reduced CD206+ expression yet are enriched for other tissue repair-associated markers. scRNAseq analysis as described in Fig. 7. (**A, B**) Violin plots further define the macrophage populations by showing the differential gene expression of alternatively activated (M2) macrophage markers (*Arg1, Chil3, Mrc1*) and classically activated (M1) macrophage markers (*Nos2, IL6, IL1b, and Ifng)* within (a) 0, 1, 2, 5 combined and (b) the individual clusters. (**C**) Volcano plots represent the differential gene expression of TGM to PBS vehicle control (clusters 0, 2, and 5). Horizontal dotted lines represent significant fold changes of expression for each treatment group. Vertical dotted lines represent a significant p-value (<0.05). Red dots represent genes significantly upregulated with the two treatment groups.

Other markers associated with the M2 macrophage phenotype, including *C1qa, Cc18*, and *Folr2* (*71*), were also found to be different between wound biopsies treated with TGM and the PBS vehicle control. *C1qa* was significantly increased in clusters 0 along with *Arg1* and *Chil3*, as well as clusters 1 and 4, while *Cc18* was decreased in clusters 0 and 2 of TGM-treated wounds (Fig. 7b, Fig. S6b). The gene *Flolr2*, which is used to identify M2 macrophages as well as tissue-resident macrophages (*71, 78, 79*), was similarly expressed between all macrophage-associated clusters except for cluster 2, where it was significantly decreased in TGM-treated wounds (Fig S6b). Additionally, *PPARγ* has been used to identify the metabolic activity denoting the M1 transition to an M2 macrophage phenotype (*80*). This gene has been observed in early monocytes in tissue repair environments, appearing during the transition to the anti-inflammatory state (*80*). Notably, this marker was significantly reduced in clusters 0, 1, and 2 in TGM-treated wounds (Fig. 8c, Fig. S6c). By contrast, markers typically associated with classical inflammatory macrophages (*IFNγ, IL-6, Nos2*) were minimally expressed in all clusters 1 through 5, with the exception of *IL-1β* which was amplified particularly in cluster 2 (Fig. 8a, b).

Genes associated with skin regeneration, fibronectin (*Fn1*) (*76, 81*), and *Ecm1* (*82, 83*), were also notably different between treatment groups and clusters. The gene, *Fn1*, which is also a marker used for M2 identification (*76*), was markedly increased in the TGM-treated macrophage-associated clusters of TGM-treated wounds, while *Ecm1* was observed at an increased fold within cluster 0 of the TGM-treated group (Fig. 8a, b, c) (*76*). This suggests that while the macrophage populations expressed in clusters 0 through 2 were generally similar in both treatment groups, macrophages within TGM-treated wounds were more specifically associated with wound healing mechanisms than their PBS vehicle control counterparts. These findings are thus consistent with our FACS analyses showing reduced CD206 expression in TGM-treated wounds, but further show marked increases in other M2 markers, many of which play important roles in tissue repair. The data shows that TGM alters the macrophage landscape of the tissue during the wound healing response with regard to the gene expression and differentiation trajectory. Taken together, these analyses suggest that macrophages induced following TGM treatment generally show a reduced expression of CD206, but upregulation of many markers associated with M2 macrophages and the wound healing process.

## DISCUSSION

In this study, we have developed a novel therapy for the treatment of cutaneous wounds that favors regenerative wound healing over fibrosis and scarring. We administered the recently identified recombinant TGM molecule, originally isolated from helminth ES products, immediately underneath a protective Tegaderm bandage. Our findings showed enhancement of wound enclosure, re-epithelialization, collagen crosslinking, and hair follicle regeneration. Our studies further showed modulation of immune cell composition and activation at the tissue repair interface, including reduced expression of CD206 by macrophage populations expressing markers important in wound healing and characteristic of M2 macrophage activation.

Cutaneous wound healing is a complex process that mitigates tissue damage and restores the integrity of damaged barrier surfaces. Rapid wound closure following injury is essential in reestablishing a protective barrier against infection and further injury (*4, 50*). However, it can also favor pro-fibrotic development over pro-regenerative mechanisms. Fibrosis and associated scarring can generate many long-term harmful effects that may potentially be avoided with improved treatments. Recent studies suggest that the later stages of tissue repair may be sufficiently plastic that a pro-fibrotic or pro-regenerative process may be favored depending upon the cytokine and cellular environment (*45-47*).

The maturation phase of wound healing is associated with key characteristics that define if the healing process favors a pro-fibrotic pathway associated with scarring or a more favorable pro-regenerative pathway (*45-47*). Scar tissue, while effective in closing the wound, is also defined by the appearance of tight parallel collagen bundle alignment. This morphological development reduces the flexibility of the skin, impacting the normal function (*1, 10, 53*). However, pro-regenerative skin tissue exhibits a cross-linked basket-weave collagen orientation providing greater flexibility to the tissue that resembles the original tissue (*54*). Furthermore, studies have shown that rapid and effective wound healing is associated with epithelial stem cells. Many of the stem cells are located within the hair follicles that are involved with the re-epithelialization of the tissue. In pro-fibrotic tissue, the lack of vessels and required growth factors impede the neogenesis of hair follicles within the new skin (*3, 46, 54*). As hair follicles contribute to the re-epithelization process, pro-regenerative skin is associated with the presence of follicles in the newly formed tissue. Our results, based upon these morphological features, show that the application of TGM is enhancing the pro-regenerative process of wound healing compared to the pro-fibrotic-like wound repair observed in the PBS vehicle control group.

Our findings indicate that enhanced wound closure mediated by TGM is, in fact, through the TGF-β receptor binding activity as demonstrated with the use of recombinant TGM truncated variants. Only the truncated TGM D1-D3, which contains the TGF-β receptor binding domains, and not the TGM D4-D5 construct, led to the enhanced wound closure observed with full-length TGM D1-D5. Following injury, many cells release TGF-β, including epithelial cells, platelets, and neutrophils (*1, 11, 32*). TGF-β has been shown to act as a chemotactic factor for macrophage recruitment and differentiation during the early pro-inflammatory phase (*62, 84*). In later stages of wound healing, TGF-β is involved in transitioning the environment away from pro-inflammatory to initiate wound healing mechanisms (*32, 62, 84*). During this phase, this cytokine is involved with the initiation of re-epithelialization, angiogenesis, deposition of matrix components, and fibroblast activation, which together mediate granulation, wound closure, and subsequent tissue maturation (*62, 84*). However, compared to non-scarred tissue, higher levels of TGF-β have been identified in scarred tissue (*85*). In wound healing, this presence of TGF-β and its associated activities are also involved with scar development (*7, 31*). Additionally, a higher expression of TGF-β is found in dysregulated fibrotic pathology such as keloid formation (*7, 85*). Interestingly, our studies suggest that TGM does not favor fibrosis and associated scarring, consistent with previous *in vitro* studies showing that administration of TGM to fibroblasts did not enhance αSMA expression to the same degree as TGFβ (*23*). Indeed, our studies showed reduced αSMA expression in skin biopsies treated with TGM compared to treatment with the vehicle control.

TGM has many characteristics that may make it a viable option for therapeutic use. While sharing the capacity to specifically bind the TGF-βR complex, TGM is structurally distinct from TGF-β and, in fact, shares more similarities with the Complement Control Protein family (*24, 63*). Furthermore, possibly due to its large size, it can bind TGF-βRI and TGF-βRII sites that are well-separated and not directly adjacent to one another as in the TGF-β receptor complex. TGM is also more bioactive than TGF-β (*63*). Additionally, and perhaps more significantly for therapeutic use, TGM does not require proteolytic cleavage for activity, and it is more stable and easier to modify for pharmacological use. Thus, while it is currently referred to as a mimic, the known activities of TGM are similar to but also distinct from mammalian TGF-β. It should also be noted that TGM may contain other as yet unidentified TGFβR-independent activities. However, our findings show a critical role for the TGFβ receptor-binding domains and further indicate the specificity of the corresponding domains for mediating wound healing activities.

A significant finding in our study was that TGM appears to reduce the presence of prototypic CD206+ M2 macrophages, confirmed through both flow cytometric and scRNAseq analysis. These observations could suggest that there is a greater number of another macrophage population reducing the proportion of CD206+ cells found within the total number of macrophages or, based upon the macrophage clusters analysis, that the CD206+ macrophages are reduced in TGM-treated wounds. Macrophages are critical to the wound healing process and are involved in the release of many pro-inflammatory and wound healing cytokines depending upon the stage of tissue repair. Previous studies have observed that in scarred versus non-scarred tissue, there are noted differences in the maturity and activation of various innate cells involved with the wound healing process (*3, 86*). The CD206+ cell subset is known to be significantly increased during wound healing. However, it is also linked to detrimental outcomes, as previous studies have shown that a high frequency of M2 macrophages in the wound bed is associated with enhanced fibrotic development (*66, 87*). However, current research favors an expanded definition of M2 macrophages (*88, 89*), inferring that CD206 may not be the only marker to define this cell population. Our results confirm this recognition of various M2 subsets as, while CD206+ expression was diminished in the macrophage populations of our TGM treated compared to the PBS vehicle control, we observed enrichment of several other M2-associated markers, including *Arg1* in some population clusters and a mixture of enrichment for *Chil3* among the macrophage populations. As such, it is possible that TGM-mediated enhanced wound repair is, in part, a result of a reduction in CD206+ macrophages with corresponding enhanced expression of other wound healing factors.

Our current study observed activity primarily on recruited macrophages, but the impact of TGM upon the resident macrophages, which are also extensively involved with the wound healing process, is still unknown. Dermal macrophages phagocytose collagen to allow remodeling and thus may help provide the tissue environment required for hair follicle development (*46, 47, 54, 90*). Within the heterogeneity of the macrophage populations identified, enhanced expression of tissue-resident gene expression (*Flolr2*) was observed in the TGM-treated wounds. Future studies examining the effects of TGM on dermal macrophages may thus provide important insights into mechanisms of TGM-mediated tissue regeneration. Through the scRNAseq analysis, we also observed in the TGM-treated group an increase in macrophage expression of genes associated with skin regeneration, specifically *Fn1* and *Ecm1*. Together, these findings suggest that CD206 may not be the appropriate marker to identify macrophages mediating wound repair following TGM treatment. Instead, our results suggest the appearance of macrophage subsets following TGM treatment that, while having reduced expression of CD206, still retain many markers associated with the wound healing process.

Our findings indicate that TGM enhances the rate of wound closure and directs the maturation phase towards a pro-regenerative process that includes marked increases in hair follicle formation and extensive collagen cross-linking. We further show extensive TGM-induced reprogramming of myeloid cells. As such, these studies provide a significant framework for the potential use of a highly purified helminth ES product as a therapy to promote cutaneous wound healing.

## MATERIALS AND METHODS

### Cell culture

A 2-D cell wound healing model system was performed using a dermal fibroblast cell line, L929, and a keratinocyte cell line, HaCaT. These cells were cultured in DMEM (Fisher Scientific) supplemented with 4mM L-glutamine (Gibco), 1mM sodium pyruvate (Gibco), 10% fetal bovine serum (FBS; Gibco), and antibiotics (1% anti-anti; Gibco) at 37ºC in an atmosphere containing 5% CO_2_ as previously described (*30*). Cells were expanded at 70-80% confluency.

### Scratch Test analysis

Scratch test analysis with SMA, TSES, HES, and TGM were performed as previously described (*15, 20, 23, 30, 91, 92*). Briefly, 24 well tissue culture plates were pretreated with 0.2mg/ml of collagen from rat tail (Sigma; 1mg/ml stock diluted in 0.1M acetic acid) for 2 hours at 37ºC and 5% CO_2_. After incubation, the plates were rinsed with warm sterile PBS. Each well was seeded with 100,000 cells in 1.2 ml of culture media. The wells contained keratinocytes alone, fibroblasts alone, or both cell types at a 50:50 ratio. The cells were incubated at 37ºC and 5% CO_2_ for 24 hours to generate a confluent monolayer (∼70-80% plate coverage). Upon confluency, each well was scored with a p1000 pipette. The media was immediately removed to collect the dislodged cells and replaced with fresh cell culture media. A marker was used on the plate underside to mark the open area to aid as an hour 0 control for the analysis over time. Two photos from each well were taken on an EVOS Core Imaging System at time 0 right after the scratch and additional photos were collected at 3, 6, 9, 12, 18, and 24 hours. The online program, Tscratch, was used to measure the area of the open wound (*93*). The percentage of wound closure was calculated by the area of the wound area at a given time compared to the wound area at hour 0. Each treatment group was performed in quintuplicate.

### Preparation of TGM

Purified recombinant *Hp*-TGM was generously provided by Dr. Rick Maizels’ lab at the University of Glasgow and prepared as previously described (*23, 63*). Briefly, the lyophilized TGM recombinant protein was resuspended in sterile H_2_0 and 35% of sterile glycerol. The TGM was aliquoted into 10μl or greater volumes for storage in the -80°C.

### Skin biopsy wound healing model

C57/BL6 mice were purchased from Jackson Laboratory (stock # 000664; Bar Harbor, Maine). For most of the wound healing studies, 8-9 week old male mice were used. The mice were maintained in a pathogen-free facility at the Rutgers New Jersey Medical School Comparative Medicine Resources. The wound healing studies used a revised protocol based upon a standard procedure (*35, 41*). Briefly, two days prior to the wounding, the mice were anesthetized with a rodent cocktail containing Ketamine (80mg/kg) and Xylazine (10mg/kg) and their dorsal skin was shaved followed by a depilatory cream (Nair) to further remove hair. On the day of the surgery, the mice were again anesthetized with the ketamine/xylazine cocktail. Following a 3-step betadine/ethyl alcohol wash, two full-thickness wounds (epidermis, dermis, and subcutaneous) were induced on each side of the midline dorsal skin of the mice using a sterile 5mm biopsy punch (Integra Miltex). A sterile transparent bio-occlusive film (Tegaderm, 3M; Saint Paul, Minnesota) was applied over the wound (*34, 41*). The mice were split into two treatment groups with one group receiving a PBS vehicle control (50ul) and the other receiving TGM (500ng). A stock of 1.5% carboxymethylcellulose in sterile PBS was made. TGM was diluted in this vehicle to generate a 500ng of TGM/50ul of solution. The 1.5% carboxymethylcellulose/PBS solution (+/- TGM) was injected through the Tegaderm and on top of each wound (50ul/wound). The mice were allowed to recover from the procedure in a warming chamber and then house separately for the duration of the study. Mice were anesthetized daily with isoflurane to image the wounds and apply an additional 50μl of the 1.5% carboxymethylcellulose/PBS vehicle control to each wound with or without 500ng of TGM at the appropriate time point. The Tegaderm was replaced every other day. The wounds were digitally photographed daily until the completion of the study. The images were blinded through the Blind Analysis plugin on Image J (NIH). They were then measured and analyzed through Image J. A ruler placed beneath the wound for each image was used to normalize and measure the area of each wound in the Image J program. The percentage of wound closure was calculated by the area of the wound at a given time compared to the wound area at Day 0. Multiple images were collected for each mouse wound and then averaged. On the day of harvest, the two wounds and surrounding tissue from each mouse were collected with 10mm biopsy punches (Acuderm Inc, Fischer Scientific). One wound biopsy was used for further flow cytometric and gene expression analysis. The other wound from the same mouse was halved with a scalpel for use in immunofluorescence and histological analysis (explained below). Studies using the truncated TGM variants were performed following the same protocol as the whole TGM treatment studies. All *in vivo* studies were performed in accordance with a protocol approved by the Institutional Animal Care and Use Committee (IACUC) at Rutgers University.

### Flow Cytometry

Single cell isolation and flow cytometric analysis of the skin was performed as previously described (*94, 95*). Excised skin was placed in a 0.5% Dispase II solution (diluted in PBS; Roche) and placed at 4ºC overnight. The following day, the digested skin was minced and further processed through a 37ºC incubation with 0.2% Collagenase Type 1 (Worthington Biochemical Corporation) in DMEM-F12+ 10% FBS for 1 1/2 hour. Erythrocytes were removed through cell lysing in ACK lysing buffer (Thermo Fisher Scientific). One 10mm biopsy punch gave approximately 2.0×10^6^ cells. 1 million cells were blocked with TruStain Fc Block (anti-mouse CD16/32, Biolegend). Cells were stained for 30 mins with antibodies specific to CD45-FITC (1:100, Biolegend), F4/80-PE (1:60, BD Biosciences), CD206-BV711 (1:100, Biolegend), CD64-PE-Cy7 (1:60, Biolegend), CD11b-BV395 (1:200, BD Biosciences), CD11c-BV421 (1:100, BD Biosciences), Ly6G-APC-Cy7 (1:60, BD Biosciences), IA/IE(MHCII)-PerCP/cy5.5 (1:125, BD Biosciences), CD301b-AF647 (1:100, Biolegend), Ly6C-AF700 (1:60, BD Biosciences). The stained cells were analyzed by flow cytometry.

### scRNA seq

Immune cells were obtained from 8-week-old C57/Bl6 mice purchased from Jackson Laboratory (stock # 000664). The wound biopsy was performed as previously described. TGM/PBS (with 1.5% carboxymethylcellulose) was administered daily at 500ng/50μl under the Tegaderm. Mice from each treatment group (n=3/group) were harvested on Day 3. Excised skin was placed in 0.5% Dispase II solution (Roche) and placed at 4ºC overnight. The following day, the digested skin was minced and further processed through a 37ºC incubation with 0.2% Collagenase Type 1 (Worthington Biochemical Corporation) in DMEM-F12+ 10% FBS for 1 1/2 hour. Erythrocytes were removed through cell lysing in ACK lysing buffer (Thermo Fisher Scientific). Cells were blocked with TruStain Fc Block (anti-mouse CD16/32, Biolegend). Cells from each sample were stained individually for 30 mins with a master mix containing the antibodies specific to CD45-FITC (1:100, Biolegend) and a TotalSeq Hashtags Antibody marker (Biolegend). Each of the three biological replicates that were treated with PBS vehicle control or TGM was labeled with one of three unique TotalSeq Hashtags Antibody markers (Biolegend). The samples were individually sort-purified by gating CD45+ cells and then each treatment group was pooled with an even number of cells from each sample. The two pooled samples (PBS and TGM) were then submitted to Rutgers genomics for further single-cell analysis using 10X genomics. For analysis, individual samples could be detected and further analyzed using the unique hashtag identifier. Briefly, as previously described (*95*), the counts were normalized and then compared between the two treatment groups to look at the differential gene expression using DESeq2.

### Histological analyses

A scalpel was used to cut the 10mm punch wound down the midline to obtain the center of the wound. One wound half was mounted in optimum cutting temperature (O.C.T.) compound (Tissue-Tek, Sakura). 5-12 thick μm cryosections were cut on a cryostat using the first few several slices for analysis to ensure the wound center was used. The sections were air dried for 30 mins to an hour and fixed in cold 99.5% histological grade Acetone (Sigma-Aldrich) for 15 mins and then immunostained as previously described (*95-97*). Briefly, following fixation, tissue sections were blocked with 10% Rat Serum (Abcam, Cambridge) and 1% CD16/32 Fc Blocking antibody (Biolegend, San Diego) in PBS for 1 hour. They were then stained overnight at 4ºC with the following antibodies: α-smooth muscle actin (αSMA for myofibroblast, AF488 1:100, Biolegend). Following primary staining, slides were stained with DAPI and then coverslips were applied using Prolong Gold Antifade Reagent (Life Technologies). Images were obtained through the Leica fluorescent microscope. Control images were used to normalize the florescence intensity of the photos taken for each sample.

The other half of the cut skin was used for H&E and picrosirius red analysis. The skin was formalin-fixed and then paraffin-embedded. 5μm slides were used to stain for H&E and picrosirius red. The first several sections were again used to obtain the wound center for analysis. Wound and wound edge thickness were measured through ImageJ software. Collagen percentage was also analyzed using the MRI Fibrosis Tool plugin through ImageJ (*56*).

### Protein analysis of serous discharge

For protein analysis of the serous discharge, 50μl of sterile PBS was added to the exudate of each wound, prior to removing the Tegaderm, and then taken up. The additional liquid improved the ability to perform a full collection of the exudate. Since the volume from each mouse was small, each treatment group was pooled providing around 250μl of solution from each treatment. The samples were spun to collect the cells. The supernatant was replaced by an additional 1 ml of sterile PBS. The protein concentration was obtained through a Nanodrop. The protein was concentrated using a 0.5 Centrifugal Filter Unit (Amicon) and then run on a Precast Protein Gel (Bio-Rad, 456-1103) (*98, 99*). Each well was loaded with 30μl of a solution containing 25ul of the protein and Pierce 5X Lane Marker Sample (ThermoFisher Scientific, Waltham). Three half dilutions of the protein, both control and TGM treated, were made, denatured in 5% of 2-Mercaptoethanol, and then loaded into the gel which was run at 100V in a 1X Tris-Glycine buffer. The gel was fixed in 50% Methanol and 10% of Acetic Acid, stained overnight in Coomassie Blue Stain, and then de-stained the next day (*99, 100*). The prepped gel was brought to the Center for Advanced Proteomics Research (Rutgers Biomedical and Health Science New Jersey Medical School) for further protein analysis by LC-MS/MS.

### Bone marrow-derived macrophage isolation and culture

Bone marrow-derived monocytes were isolated from the femur and tibia of 8-week-old C57/Bl6 mice (n=3) as previously described (*101-103*). Erythrocytes were removed through cell lysing in ACK lysing buffer (Thermo Fisher Scientific) and resuspended in DMEM:F12 LCM lymphocyte growth media, which was obtained from L929 conditioned culture media and prepared and treated for *in vitro* differentiation as previously described (*102, 104*). Briefly, cells were plated at 500,000 cells/well in a 6-well plate. On day 6, the matured macrophages were stimulated for 16 hours. All cells received MCSF (50 ng/ml, R&D Systems). Both recombinant LPS (100ng/ml) and recombinant IFNy (10 ng/ml, Gibco) were provided to cells for differentiation to classical macrophages. Recombinant IL-4 (25ng/ml, PeproTech) and IL-13 (25ng/ml, R&D Systems) were provided to cells for alternatively activated macrophage differentiation. After incubation, the macrophages were trypsinized and scraped out of the well, centrifuged at 200 x g for 10 minutes, and then resuspended in fresh lymphocyte media. The cells were processed for flow cytometry as described in the earlier methods. Cells were stained for 30 mins with antibodies specific to CD45-FITC (1:100, Biolegend), F4/80-PE (1:60, BD Biosciences), CD206-BV711 (1:100, Biolegend), CD64-PE-Cy7 (1:60, Biolegend), CD11b-BV395 (1:100, BD Biosciences), CD11c-BV421 (1:100, BD Biosciences), Ly6C-AF700 (1:60, BD Biosciences). The stained cells were analyzed by flow cytometry.

### Statistical analyses

Data were analyzed with Prism 9 (GraphPad) and reported as mean +/- standard error. One-way ANOVA and protected T-tests were used to assess the difference between multiple groups at each time point. Comparisons between two groups were analyzed through a Student’s T-test. Each *in vivo* timepoint study or antibody study was replicated two or more times. A P value of <0.05 was considered significant.

## Funding

National Institute of Health T32 Training Grant 5T32AI125185-04 (WCG)

National Institute of Health R01 Award 1R01AI131634 (WCG)

Wellcome Trust through an Investigator Award to RMM 106122 and 219530 (RMM)

Wellcome Trust core-funded Wellcome Centre for Integrative Parasitology 104111 (RMM)

## Notes

### Competing Interest Statement

The authors have declared no competing interest.

