## Supplemental Figures for "A helminth mimic of TGF-β, TGM, enhances regenerative cutaneous wound healing and modulates immune cell recruitment and activation"

### Supplemental data

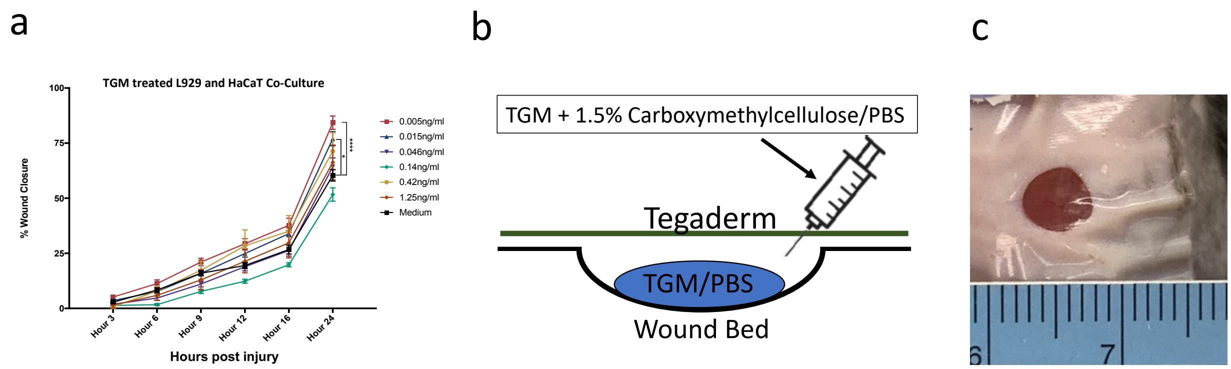


**Fig. S1. TGM accelerates cell migration in an *in vitro* scratch test.** (**A**) A 2-D *in vitro* scratch test wound model was generated with a 50:50 co-culture of L929 fibroblast and HaCaT keratinocytes to examine the wound closure and migration with the application of TGM. The area of wound remaining open, calculated by Tscratch, was used to quantify the rate of wound closure for different concentrations of TGM (0.005ng – 1.25ng/ml) compared to media alone from 0 to 24 hours. The percentage of wound closure at each time point is compared to the percentage open at hour 0 (** $p < 0.01$, *** $p < 0.001$, **** $p < 0.0001$; $n = 5$ independent wells per condition). Statistical analysis was performed using a two–way analysis of variance (ANOVA) test with Tukey's multiple comparisons for comparison between all treatment groups at each timepoint. Error bars represent mean+/- SEM. Results are representative of two or more independent experiments. (**B**) 5mm full-thickness excisional wounds were generated on the dorsal skin of C57BL/6 mice. Wounds were treated with PBS vehicle control or TGM (500ng) and covered with Tegaderm© for the duration of the study. The diagram depicts the application of TGM within a 1.5% carboxymethylcellulose vehicle administered underneath a layer of Tegaderm to replicate a topical application of the molecule to the wound. (**C**) A representative image of a wound covered by Tegaderm on day 1. The ruler was used for each image as a control for measuring the area of the open wound.

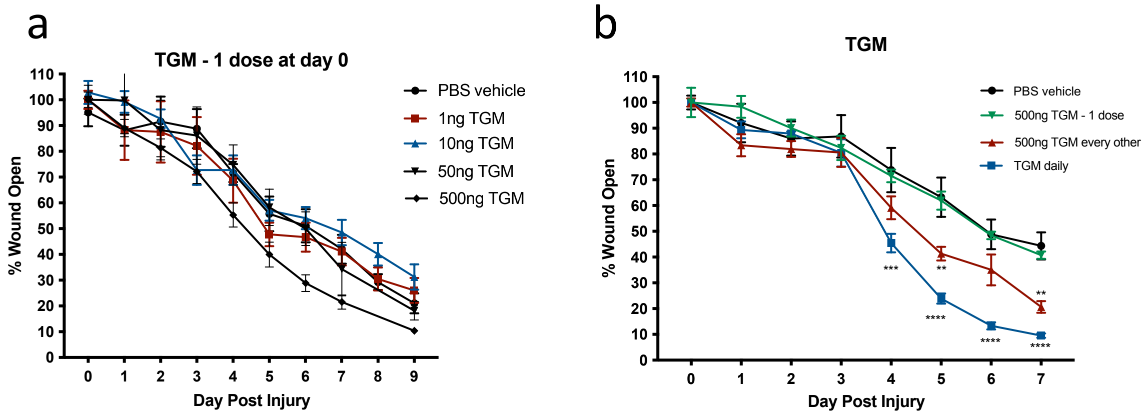


**Fig. S2. TGM given at one dose at day 0 in each wound did not lead to enhanced wound closure in an *in vivo* biopsy model.** (**A**) 5mm full-thickness excisional wounds were generated on the dorsal skin of C57BL/6 mice. Wounds were treated with PBS vehicle control or various concentrations of TGM (1ng – 500ng) and covered with Tegaderm© for the duration of the study. Wound size analysis was performed on the gross images obtained at each time point during the course of treatment. Treatments were given once on day 0 while the dressing was changed every other day. (**B**) The results from a wound biopsy study comparing PBS vehicle control and various TGM (500ng) dosing regimens: one day (at day 0 only), every other day (at days 0, 2, 4, and 6), and daily (given every day) and covered with Tegaderm© for the duration of the study. (A, B) Wound closure rates with the various concentrations (1ng – 500ng) or different dosing regimens (500ng) of topical TGM or PBS vehicle control over 7 or 9 days were quantified as the percentage of wound closure at each time point compared to the percentage open at Day 0. (** $p < 0.01$, *** $p < 0.001$, **** $p < 0.0001$; 5 independent wounds from each treatment were measured through blinded analysis on ImageJ). Statistical analysis was performed using a two–way analysis of variance (ANOVA) test with Tukey's multiple comparisons for comparison between all treatment groups at each timepoint. Error bars represent mean+/- SEM. Results are representative of two or more independent experiments.

a

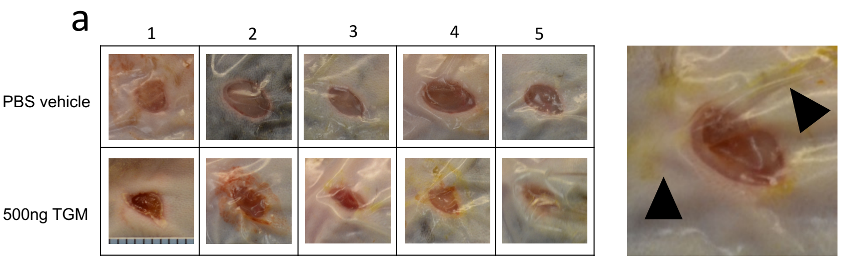


b

| Description | Accession | Coverage [%] | Spectra counts (Ctrl) | Spectra counts (Treated) | Ratio (Treated/Ctrl) |
|---|---|---|---|---|---|
| Serum amyloid A-1 protein OS=Mus musculus OX=10090 GN=S | P05366 | 54.09836066 | 1 | 9 | 3.7 |
| Basement membrane-specific heparan sulfate proteoglycan co | E9PZ16 | 7.00433493 | 5 | 20 | 3.1 |
| Retinoic acid receptor responder protein 2 OS=Mus musculus ( | A0A0N4SVV4 | 50.30674847 | 1 | 7 | 3.0 |
| GDH/6PGL endoplasmic bifunctional protein OS=Mus musculu | Q8CFX1 | 5.830164766 | 0 | 4 | 3.0 |
| Procollagen-lysine,2-oxoglutarate 5-dioxygenase 1 OS=Mus mu | Q9R0E2 | 6.043956044 | 0 | 4 | 3.0 |
| Thrombospondin-4 OS=Mus musculus OX=10090 GN=Thbs4 PE | Q9Z1T2 | 22.74143302 | 7 | 23 | 2.8 |
| Hepatocyte growth factor-like protein OS=Mus musculus OX=1 | E0CXN0 | 10.32531825 | 1 | 6 | 2.7 |
| Thrombospondin-3 OS=Mus musculus OX=10090 GN=Thbs3 PE | E9PWF0 | 4.352557127 | 1 | 6 | 2.7 |

c

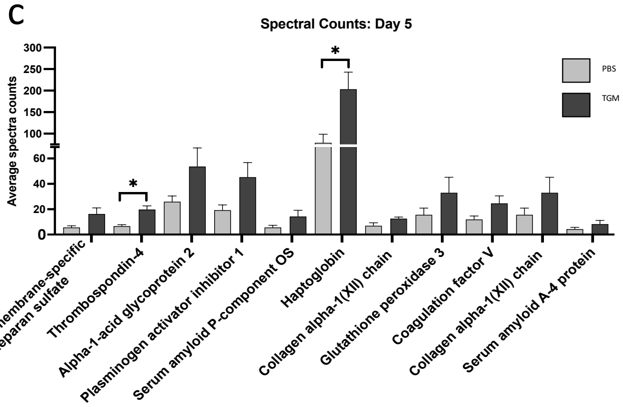


**Fig. S3. Enhanced serous secretion containing wound healing associated protein is observed at a higher frequency in the TGM-treated wounds.** (**A**) Representative wound images from day 5 of mice treated every other day topically with PBS vehicle control or TGM (500ng) demonstrate an increased quantity of discharge underneath the Tegaderm of each wound treated with TGM. Numbers (1-5) represent each mouse used in the study (n=5). The black arrows identify the discharge in a magnified image of a wound treated with 500ngs TGM on day 5. (**B**) Graphs represent raw LC-MS/MS analysis output of the protein composition from one of three repeat protein analysis studies performed on the serous discharge collected from wounds collected on day 5 after receiving PBS vehicle control and TGM every other day starting at day 0. (**C**) The bar graph represents the spectral counts of the protein content between the two groups. (B, C) Statistical analysis was performed using a t-test to compare the two treatments for each represented protein. (* $p < 0.05$) Error bars represent mean+/- SEM. Results from two or more independent determinations demonstrated similar results.

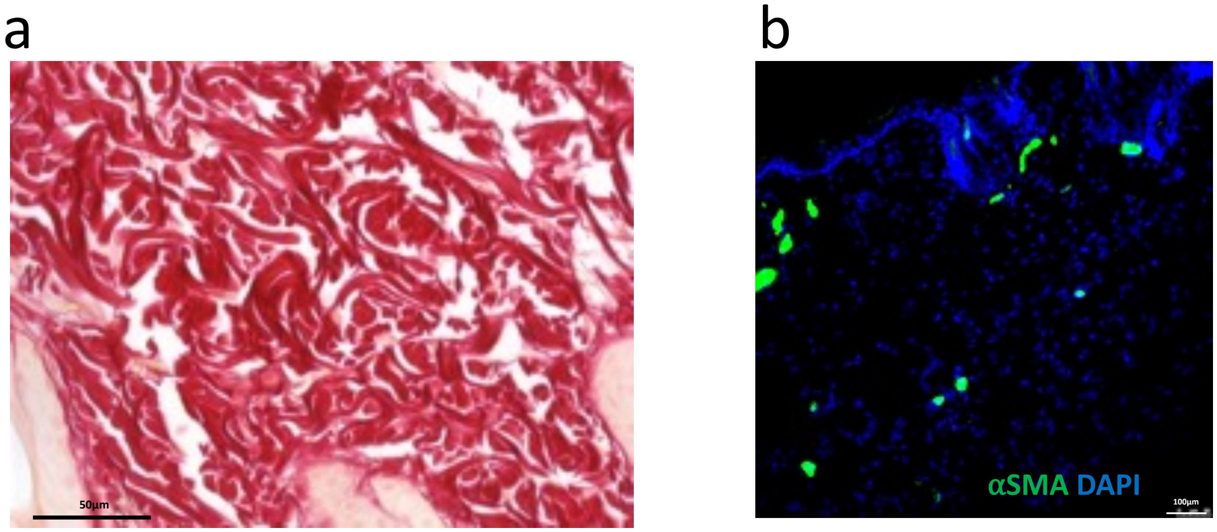


**Fig. S4. Nonwounded skin morphology.** (**A**) A representative picrosirius red-stained wound image from the unwounded skin of a mouse shows collagen with a basket-weave orientation typical of normal skin. (**B**) A representative immunofluorescent stained image of a wound bed stained with αSMA on unwounded skin with no TGM treatment; αSMA (green fluorescent signal), DAPI (blue fluorescent signal). Immunostained slides with αSMA were quantified on unwounded skin by the fluorescent intensity measured as pixel area using ImageJ. (* $p < 0.05$; 5 biologically independent samples were used per treatment for the blinded analysis on ImageJ). Results from two or more independent determinations demonstrated similar results

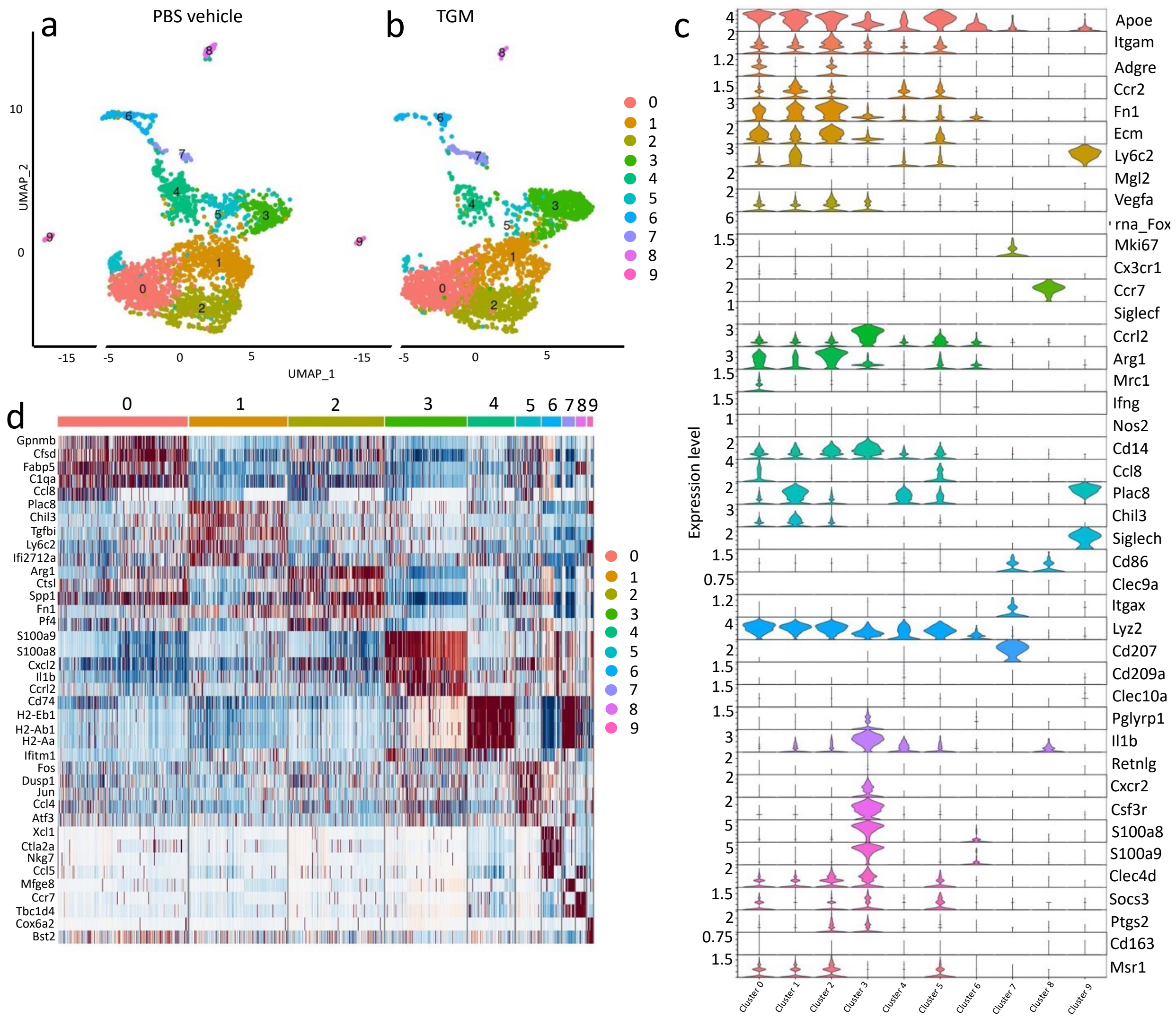


**Fig. S5. scRNAseq analysis identifies distinct clusters within the skin at day 3 post wounding.** scRNAseq analysis of CD45+ leukocytes obtained from wound bed at day 3 after skin injury. Skin wounding was performed as described in Fig. 1. A uniform manifold approximation projection (UMAP) plot of the single cells obtained from CD45+ cells purified from (**A**) PBS vehicle control and (**B**) TGM treated wounds on day 3 after skin injury. 10X genomics scRNAseq using was performed on the CD45+ purified cells treated with PBS vehicle control or TGM (3 biologically independent samples were used per treatment labeled with 3 unique TotalSeq Hashtag Antibody markers). The analysis identified 9 distinct cell clusters varying between treatment groups (b) Violin plots highlight the additional markers used to define the populations within the clusters. (**C**) A heatmap identifies the top 5 differentially expressed markers within each cluster.

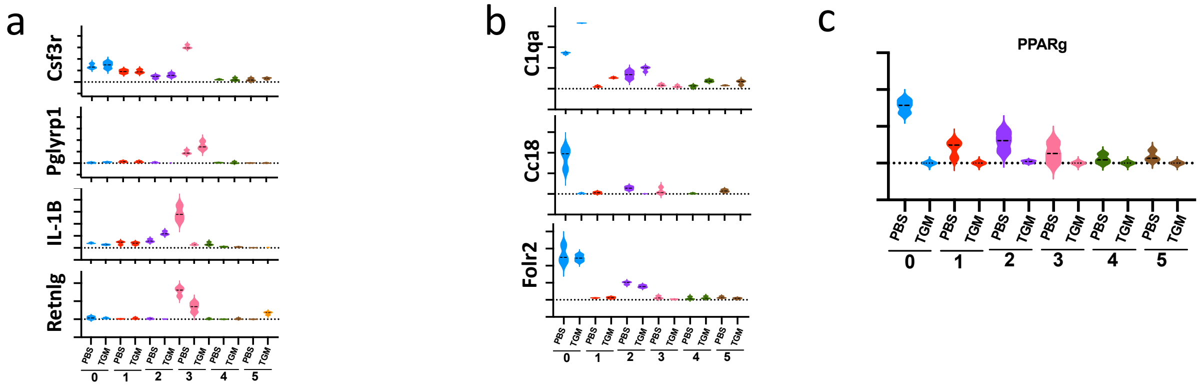


**Fig. S6. scRNAseq highlights the differential expression of neutrophils and macrophages subsets between treatment groups.** The scRNAseq analysis of CD45+ cells purified from day 3 PBS vehicle control and TGM treated wounds are displayed in violin plots which represent the differential gene expression of (**A**) neutrophil-associated markers (*Csf3r, Pglyrp1, Il1b, Retnly*). The dotted line represents the gene expression cutoff. The dotted lines within the graphed violin plot represent the mean. (**B**, **C**) Violin plots represent the differential gene expression of additional (B) M2 macrophage markers (*C1qa, Ccl8, and Folr2*) as well as the (C) M2 associated metabolic marker *Ppary*. The dotted line represents the gene expression cutoff while the dotted lines within the graphed violin plot represent the mean.
